# Genome-wide identification, characterization, and comparative analysis of NLR resistance genes in *Coffea spp*

**DOI:** 10.1101/2022.02.03.478062

**Authors:** Mariana de Lima Santos, Mário Lúcio Vilela de Resende, Gabriel Sérgio Costa Alves, Jose Carlos Huguet-Tapia, Márcio Fernando Ribeiro de Resende, Jeremy Todd Brawner

**Affiliations:** Laboratório de Fisiologia do Parasitismo, Faculdade de Ciências Agrárias, Departamento de Fitopatologia, Universidade Federal de Lavras, Lavras, Minas Gerais, Brazil; Laboratório de Processos Biológicos e Produtos Biotecnológicos, Instituto de Ciências Biológicas, Departamento de Biologia Celular, Universidade de Brasília, Brasília, Distrito Federal, Brazil; Institute of Food and Agricultural Sciences, Department of Plant Pathology, University of Florida, Gainesville, Florida, FL, United States; Institute of Food and Agricultural Sciences, Horticultural Sciences Department, University of Florida, Gainesville, Florida, FL, United States

**Keywords:** resistance genes, Coffea, nucleotide-binding site leucine-rich repeat, Genome-wide, NLR-annotator

## Abstract

The largest family of disease resistance genes in plants are nucleotide-binding site leucine-rich repeat genes (NLRs). The products of these genes are responsible for recognizing avirulence proteins (Avr) of phytopathogens and triggering specific defense responses. Identifying NLRs in plant genomes with standard gene annotation software is challenging due to their multidomain nature, sequence diversity, and clustered genomic distribution. We present the results of a genome-wide scan and comparative analysis of NLR loci in three coffee species (*Coffea canephora, Coffea eugenioides* and their interspecific hybrid *Coffea arabica*). A total of 1311 non-redundant NLR loci were identified in *C. arabica*, 927 in *C. canephora*, and 1079 in *C. eugenioides*, of which 809, 562, and 695 are complete loci, respectively. The NLR-annotator tool used in this study showed extremely high sensitivities and specificities (over 99%) and increased the detection of putative NLRs in the reference coffee genomes. The NLRs loci in coffee are distributed among all chromosomes and are organized mostly in clusters. The *C. arabica* genome presented a smaller number of NLR loci when compared to the sum of the parental genomes (*C. canephora*, and *C. eugenioides*). There are orthologous NLRs (orthogroups) shared between coffee, tomato, potato, and reference NLRs and those that are shared only among coffee species, which provides clues about the functionality and evolutionary history of these orthogroups. Phylogenetic analysis demonstrated orthologous NLRs shared between *C. arabica* and the parental genomes and those that were possibly lost. The NLR family members in coffee are subdivided into two main groups: TIR-NLR (TNL) and non-TNL. The non-TNLs seem to represent a repertoire of resistance genes that are important in coffee. These results will support functional studies and contribute to a more precise use of these genes for breeding disease-resistant coffee cultivars.

## 1 Introduction

Coffee is a globally important agricultural commodity that plays a significant economic role in producing and consuming countries (Krishnan, 2017). The genus *Coffea* consists of more than 100 botanical species (Davis et al., 2006), however, the most cultivated species are *Coffea canephora* and *Coffea arabica. C. canephora* is diploid (2n = 2x = 22 chromosomes) (Denoeud et al., 2014), while *C. arabica* is a allotetraploid (2n = 4x = 44 chromosomes) (Tran et al., 2018) originated from natural hybridization between *C. canephora* and *C. eugenidoides* (Lashermes et al., 1999; Bawin et al., 2020). Among the more than 50 coffee-producing countries, Brazil, Vietnam, Colombia, and Indonesia are major producers, with Brazil being the largest producer by volume. Currently, coffee diseases are the main factor affecting productivity (Cerda et al., 2017). Examples of diseases associated with coffee include cercosporiosis (*Cercospora coffeicola*), bacterial blight (*Pseudomonas syringae* pv. *Garcae*), anthracnose (*Colletotrichum coffeanum*), root-knot nematodes (*Meloidogyne spp*.), coffee berry disease – CBD (*Colletotrichum kahawae*), and coffee leaf rust – CLR (*Hemileia vastatrix*) (Cabral et al., 2016; Krishnan, 2017). CLR is one of the most devastating diseases found in coffee and is present in all regions of the world where coffee is grown (McCook and Vandermeer, 2015; Cabral et al., 2016). Currently, 95% of *C. arabica* varieties cultivated in Brazil are susceptible to CLR due to the emergence of variants of the pathogen (Cabral et al., 2016). Given the increasing problem of plant pathogens in coffee production, a greater understanding of the set of receptors regulating the plant immune system of coffee is needed.

Throughout evolution, plants have developed sophisticated systems to defend themselves from pathogens. The plant immune system involves both broad-spectrum and specific recognition of pathogens. Broad-spectrum recognition is related to the detection of pathogen associated molecular patterns (PAMP), such as fungal chitin or bacterial flagella, by pattern recognition receptors (PRR) that are anchored to the plasma membrane and trigger the PAMP-triggered immunity (PTI) (Boutrot and Zipfel, 2017). Specific recognition, on the other hand, primarily involves receptors encoded by resistance genes (R genes) that detect the presence of pathogen effector proteins and trigger effector-triggered immunity (ETI) (Jones and Dangl, 2006). Both types of recognition occur dynamically and continuously, converging into signaling pathways that activate essential mechanisms for downstream responses to pathogen recognition (Lu and Tsuda, 2021; Yuan et al., 2021).

The R genes have been extensively studied in several crops to facilitate their greater use in plant breeding (Jupe et al., 2013; Wan et al., 2013; Lozano et al., 2015; Inturrisi et al., 2020; Steuernagel et al., 2020). The protein products of these genes recognize directly or indirectly effector proteins that are secreted by pathogens (Kourelis and Van Der Hoorn, 2018) and trigger a series of signaling steps that lead to the hypersensitive response (HR) that activates cell death and potentially leads to systemic acquired resistance (SAR) (Kachroo and Robin, 2013; Jones et al., 2016). The largest and most diverse group of R genes found in plants belong to the nucleotide-binding site leucine-rich repeat family (NLR or NBS-LRR) (Jones et al., 2016). The proteins encoded by these genes are typically modular, and many have a variable N-terminal domain-containing Toll/interleucina-1 (TIR) or coiled-coil (CC) receptors. As well, the nucleotide-binding domain (NB-ARC or NBS) is a canonical feature of NRLs, that is shared with human apoptotic protease-activating factor-1 (*APAF-1*) and *Caenorhabditis elegans* death-4 (*CED-4*) proteins. A C-terminal region comprising a variable number of leucine-rich repeats (LRRs) is another common feature in NLR genes (Jones et al., 2016; Shao et al., 2019). The NB-ARC domain is highly conserved and is involved in the active and inactive state of the NLS protein (Bonardi et al., 2012; Jones et al., 2016). This domain presents motifs that are characteristic of the ATPase family, such as p-loop, kinase 2, and the RNBS (Resistance Nucleotide Binding Site) A, RNBS-C, and RNBS-D motifs (Van Ghelder et al., 2019). Mutations in specific residues within these motifs may cause the loss of protein function or self-activation and interfere with the regulation or activation of defense mechanisms. (Monteiro and Nishimura, 2018; Bezerra-Neto et al., 2020).

The domains described above provide a more refined classification of NLRs proteins, which may be characterized into two main groups: TNLs (TIR-NLRs) or non-TNL (which include CNLs - CC-NLRs). The truncation of a single domain, such as LRR (CN or TN), TIR or CC (NL), in both C and N terminal domains (N), may also be used to classify NLR genes (Monteiro and Nishimura, 2018). Additionally, atypical or non-canonical integrated domains (IDs) that act as decoys and play roles in oligomerization or downstream signaling may be present, demonstrating the structural diversity of this NLR family (Kroj et al., 2016; Wang et al., 2021). The number of NLRs in plant genomes varies greatly and is often organized in tandem, which facilitates duplication, contraction, and transposition and provides a reservoir of genetic variation that allows plant evolutionary dynamics to respond to changes phytopathogen populations (Barragan and Weigel, 2021). These genes are often under selection pressure, resulting in a large number of pseudogenes and variable loci content within the same species, among species, and across plant populations (Schatz et al., 2014; Steuernagel et al., 2015; Sun et al., 2020; Hufford et al., 2021).

The knowledge of how NLRs are distributed throughout the genome and their diversity is of great interest as it may reveal new sources of resistance that may be used to develop new cultivars (Monteiro and Nishimura, 2018). The growing number of sequenced plant genomes facilitates the search for novel NLR and has led to the genome-wide analysis of NLR genes (Denoeud et al., 2014; Song et al., 2015; Scott et al., 2020; Wang et al., 2021). However, its large number, frequently clustered genomic distribution, and low expression in uninfected tissues make cataloging NLR genes challenging and often underestimates the number of NLRs in genomes (Jupe et al., 2013; Steuernagel et al., 2015, 2020). To mitigate this problem, some specific gene/loci NLR annotation pipelines have been developed to augment standard gene annotation software and improve our ability to identify and locate genes belonging to this family. Some examples of these pipelines are NBSPred (Kushwaha et al., 2016), NLGenomeSweeper (Toda et al., 2020), and NLR-annotator, a new version of NLR-parser (Steuernagel et al., 2015, 2020). The NLR-Annotator is a tool used to annotate NLR loci that use 20 highly curated motifs present in NLR proteins and does not depend on the support of transcript data (Jupe et al., 2012; Steuernagel et al., 2020). Since it was published, this tool has been applied in several studies to prospect and annotate R genes in recently sequenced genomes (Muliyar et al., 2020; Read et al., 2020; Scott et al., 2020), to check the completeness of previous annotations (Muliyar et al., 2020), and for studies of the resistance-related locus (Jost et al., 2020).

The genome of *C. canephora* was published in 2014, which allowed the first genome-wide NLR study in coffee (Denoeud et al., 2014). In 2018, the *C. arabica* and *C. eugenioides* genomes were deposited at the NCBI, providing an essential resource for studying the structure and evolution of NLRs in arabica coffee and the contribution of the genomes that gave rise to this species. For coffee production to continue advancing in producing regions worldwide, adequate disease management is of great importance. A range of strategies must be used to control the main phytosanitary problems associated with coffee production. Using these genomic resources is essential for informing breeding strategies aimed at developing resistance to disease in coffee. Given the above, this study aimed to: (i) identify NLR loci in *C. arabica, C. canephora*, and *C. eugenioides* genomes using the NLR-annotator tool and discuss the improvements in annotation derived from the use of a specific pipeline for NLR genes in coffee, (ii) catalog, classify and characterize the distribution of NLRs loci in the *coffee spp*. genomes, and (iii) understand the contribution of *C. canephora* and *C. eugenioides* to the NLR repertoire of *C. arabica*.

## 2 Materials and Methods

### 2.1 Coffee Genomic Resources

Three genomes were used in this study. The *C. arabica* (Caturra red - Cara_1.0, GenBank assembly accession: GCA_003713225.1) and *C. eugenioides* (Ceug_1.0, GenBank assembly accession: GCA_003713205.1) genomes are available from the NCBI (National Center for Biotechnology Information) database (https://www.ncbi.nlm.nih.gov/) and the *C. canephora* genome is available at Coffee genome hub (https://coffee-genome.org/) (Denoeud et al., 2014). For the three species, the genome files, sets of predicted proteins, predicted genes, and GFF (General Feature Format) were used.

### 2.2 Identification of loci NLR in *Coffea* spp. genomes

The identification of NLR loci in *Coffea spp*. was accomplished by the NLR-Annotator, (Steuernagel et al., 2020) using the default parameters. The tool uses combinations of short motifs of 15 to 50 amino acids to classify a genomic locus as an NLR. These motifs were defined using domains of known NLR proteins used as a training set in a study carried out by Jupe et al. 2012 (Supplementary Table 1).

In summary, the pipeline is divided into three steps: 1) dissection of genomic input sequence into 20-kb fragments overlapping by 5 kb; 2) translating each fragment into all six reading frames and searching for the motifs associated with NLR by NLR-Parser to create an xml-based report file. The NLR-Parser searches for combinations of doublets or triplets of motifs and records their nucleotide positions, disregarding motifs that occur randomly. Finally in step 3, the NLR-Annotator uses the xml file as input, integrates data from all fragments, evaluates positions and combinations of motifs. In this step, the NB-ARC motifs are used as the principal seed to annotate NLR loci, generate output files (.gff, .bed - Browser Extensible Data, .txt and file of the NB-ARC motifs as multiple alignments to complete loci) based on coordinates and orientation the initial input genomic sequence (Steuernagel et al., 2020).

Each section of the genomic sequence associated with a single NLR is called an ‘NLR locus’ and this refers to an NB-ARC domain (or associated motif) followed or not by one or more leucine-rich repeats (LRRs). From the sets of motifs that are identified, these loci are classified as complete (containing the P-loop, at least three consecutive NB-ARC motifs, and at least one LRR), complete (pseudogenes), partial and partial (pseudogenes) (Supplementary Table 1). Therefore, the NLR-Annotator identifies the NLR loci that are either active genes or pseudogenes. The number of NLR loci and their classification is described in the output file.txt (Steuernagel et al., 2020 and https://github.com/steuernb/NLR-Annotator).

### 2.3 Validation of the NLR-annotator sensitivity and specificity in coffee genomes

To validate the sensitivity and specificity of the NLR-annotator in the coffee genomes, we initially classified the protein sequences of the three genomes using PfamScan (https://pfam.xfam.org/) version 1.5 with an e-value of less than 1E-5 and models from Pfam-A. Subsequently, proteins that had the NB-ARC domain (PF00931) were filtered, and from this process, we obtained the ID of the genes corresponding to each protein. With the list of gene model IDs of the NLR family, it was then possible to filter the GFF files and obtain the positions of the genes that had already been annotated in each genome.

We identified overlapping intervals to compare the NLR loci detected by NLR-Annotator and the NLR genes that had already been annotated in the genomes. We used the information from .gff files from both annotations for an overlay analysis using bedtools intersect (version 2.29.2). An overlap was only considered if both, the locus, and gene, were on the same strand. This analysis made it possible to distinguish the loci identified by NLR-annotator that were or were not overlapping with the gene models from the reference genomes.

For NLR genes already annotated in the genomes and not identified by NLR-annotator, a search for motifs by NLR-Parser was performed to obtain the xml and txt output (options -c and -o) as well the detection of conserved domains using the NCBI Conserved Domain Database (https://www.ncbi.nlm.nih.gov/Structure/cdd/wrpsb.cgi) for nucleotides sequences. Standard parameters were used for the conserved domains analysis, except for the threshold (E-value), which was set to 1E-5. The Graphical summary was set to provide a concise view of the results. To characterize the NLR loci only found by NLR-annotator and to make sure they were homologous with NLRs already annotated in plants, we aligned the sequences for these loci with NCBI’s non-redundant protein database (nr) (https://blast.ncbi.nlm.nih.gov) using BLASTx (BLAST - version 2.10.1 with the max_target_seqs option set to 5). For loci that did not have homology with NLRs proteins, a conserved domain analysis was also performed as previously described.

The sensitivity of the pipeline was calculated as the ratio of the number of loci identified by NLR-annotator (including motifs detected by NLR-Parser in the second step of the pipeline) to the number of NLRs genes already annotated in the genomes. The specificity was calculated as the ratio of the number of loci identified by NLR-annotator that are related to NLRs genes or have characteristic domains of that family to the total number of loci identified. Characteristic domains were defined as domains overlapping with the annotations already described in the studied genomes, homology with NLR proteins by BLASTx or NB-ARC domains identified with conserved domains analysis.

### 2.4 Distribution of NLR loci in Coffee’s chromosomes

In order to visualize the distribution of NLR loci on chromosomes of the three analyzed coffee species, the annotation files from NLR annotator (.txt) were used to extract the genomic position and classifications of the loci. The chromosome size information in Mb was obtained from the NCBI (for *C. arabica* and *C. eugenioides*) and Coffee genome hub (for *C. canephora*) and the visualization was created using the R software with the chromoMap package (Anand and Lopez, 2020). ChromoMap, divides the chromosomes as a continuous composition of loci. Each locus, consist of a specific genomic range determined algorithmically based on chromosome length and then the annotations are inserted. The detailed annotation information on each locus NLRs (complete, complete pseudogene, partial and partial pseudogene) is displayed in an HTML file.

### 2.5 Prediction of genes in the complete loci found only by the NLR-annotator

Gene prediction was performed using the AUGUSTUS program version 3.3.3 (http://bioinf.uni-greifswald.de/augustus/) (Stanke et al., 2006) using gene models from *Solanum lycopersicum* and allowing for the prediction of only complete genes.

### 2.6 Orthologous groups and Phylogenetic Analyses

The complete loci identified in the coffee genomes by the NLR-annotator, being those loci that overlap with gene models of the reference genomes and loci that were annotated by AUGUSTUS as putative genes were the focus of ortholog and phylogenetic analysis. In order to make a comparison with the set of coffee NLRs, 326 NLR loci identified in tomato (*Solanum licopersicum* - Heinz 1706) by Andolfo et al., 2014, 755 loci identified in potato (*Solanum tuberosum*) by Jupe et al., 2013, 67 NLR reference genes (functionally characterized protein) obtained from The Plant Resistance Genes database - PRGDB (http://prgdb.org/prgdb/, S2 table) (Osuna-Cruz et al., 2018) and the *CED-4* gene from *Caenorhabditis elegans* (outgroup) were also added. All these sequences were classified according to the rules of motifs established by the NLR-annotator and only those considered as complete NLR were used for these analyses. This criterion was used to standardize the methodology for classifying loci as complete or not.

The amino acid sequences of the NB-ARC domain were extracted from the set of complete NLR loci for the 5 species (*C. arabica, C. canephora, C. eugenioides, S. licopersicum* and *S. tuberosum*) along with the reference genes by NLR -annotator (parameter -a). All NB-ARC domain of these complete loci (hereafter called NLRs) were compared with each other using BLASTP, all-by-all (E value <1e-10). The markov clustering algorithm was performed with inflation value of 1.5 and then NLRs in the same cluster were classified as orthologous subgroups by OrthoMCL version 1.4 (standard parameters) (Li et al., 2003). In order to analyze and visualize the number of orthogroups shared between the species and the ones that are unique to a single species, we used the UpSetR package in R (Conway et al., 2017).

The NLRs clustered into single-copy orthogroups (orthogroups that have one copy of each NLR present once in each of the 5 genomes or reference NLRs) by OrthoMCL were used to construct a phylogenetic tree. The sequences were aligned using MAFFT version 6.903 (Katoh et al., 2002), with the --auto parameter to select the best alignment strategy. The tree was inferred using RAxML version 8.2.10 (Stamatakis, 2014) with the PROTGAMMAAUTO model (the JTT model was selected as having the highest likelihood) with 100 bootstrap replicates. A second phylogenetic tree classifying the coffee NLRs was also constructed with the above-mentioned parameters using the entire set of complete NLRs. Coffee NLRs were classified in the tree using the previous classification describing 67 reference NLRs (Supplementary Table 2) and the tomato and potato NLRs (Jupe et al., 2012, 2013; Andolfo et al., 2014). The trees were visualized and edited using the Interactive Tree of Life (iTOL) tool (Letunic and Bork, 2021).

## 3 Results

### 3.1 NLRs Identification, validation of the sensitivity and specificity of NLR-annotator in coffee genomes

A total of 1318 loci were identified for *C. arabica*, 932 for *C. canephora*, and 1081 for *C. eugenioides* (Supplementary Table 3). In each species, we identified some loci that are in the same position but were separated by the NLR-annotator as two distinct NLRs. We found 7, 5, and 2 repeated loci in the aforementioned species, respectively (Supplementary Table 3 highlighted in blue). Considering these repeated loci when counting the number and distribution of the NLRs loci on the chromosomes, the most complete classification was considered. After the identification of these regions, it was found that there were 1311 non-redundant loci for *C. arabica* (627 from the *C. canephora* subgenome - CaC, 650 from the *C. eugenioides* subgenome - CaE and 34 Unassigned - Un), 927 for *C. canephora* (559 mapped on chromosomes and 367 Un) and 1079 for *C. eugenioides* (944 mapped on chromosomes and 135 Un). The number of complete loci found in each species was 809 (*C. arabica*), 562 (*C. canehora*), and 695 (*C.eugenioides*).

To examine whether there was a consensus between the gene models for NLRs that have previously been annotated in the genomes and loci identified by NLR-annotator, an overlap analysis was performed. PfamScan analyses were conducted on the set of predicted proteins and the subsequent selection of NLR proteins annotated in each genome showed that 1015, 709, and 869 genes encoded proteins (including isoforms) containing the NB-ARC domain in the *C. arabica, C. canephora*, and *C. eugenioides* genomes, respectively (Figure 1, Supplementary Table 4). The overlap between the genomic positions of these genes and the positions of loci from NLR-annotator showed that of 1311, 927, and 1079 loci identified by NLR-annotator for *C. arabica, C. canephora*, and *C.eugenioides* respectively, 1013 (99,80%), 687 (96,90%), and 857 (98,62%) overlap with the genes already annotated in the reference genomes. A total of 298, 240 and 222 non overlapping NLRs were found, respectively (Figure 1, Table 1). We also noticed that there are cases in which more than one NLR loci overlapped with a single NLR gene, and the opposite was also found in all three genomes. A Venn diagram representing these data is shown in figure 1 as the intersection and data are highlighted in Supplementary Table 5.

**Figure 1.**
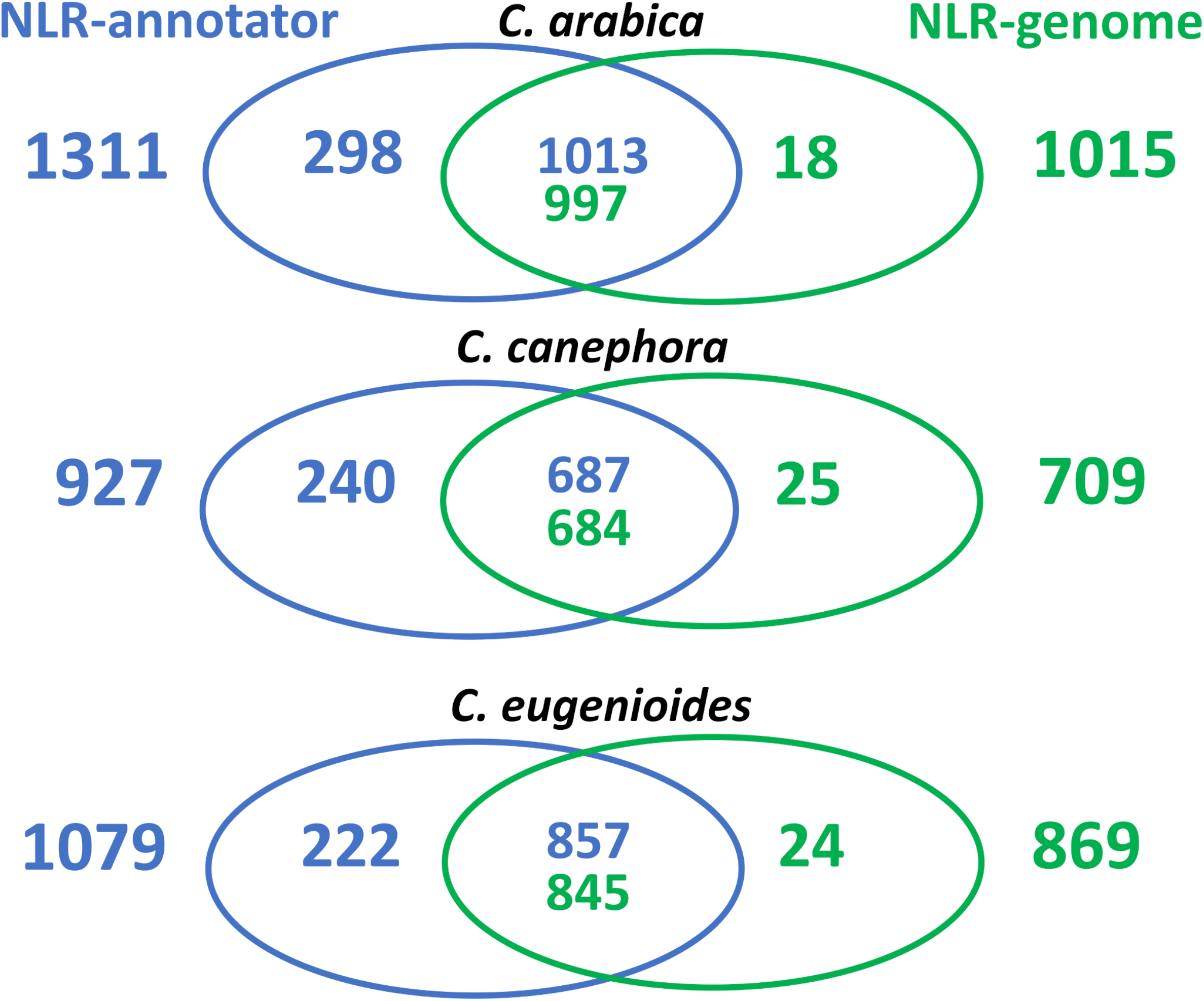
Venn diagrams representing the overlap between the loci from NLR-annotator and NLR genes annotated in the *C. arabica, C. canephora*, and *C. eugenioides* reference genomes. The colors represent the origin of the annotation, with blue indicating those annotated by NLR-annotator and green indicating those found in the reference genome. The intersection refers to the overlaps that occurred once or more than once.

**Table 1.** Loci identified using NLR-annotator that did not overlap with annotations of NLR genes from coffee reference genomes.

| Species | Complete | Complete<br>(pseudogene) | Partial | Partial<br>(pseudogene) | Total |
| --- | --- | --- | --- | --- | --- |
| *C. arabica* | 70 | 90 | 65 | 73 | 298 |
| *C. canephora* | 67 | 56 | 73 | 44 | 240 |
| *C. eugenioides* | 71 | 65 | 37 | 49 | 222 |

The overlap analysis also made it possible to identify genes annotated in the reference genomes that did not overlap with any locus from NLR-annotator. To examine these genes, an NLR-parser analysis, with options -c (file.xml) and -o (file.txt), was performed on this set. Among the genes not identified by NLR-annotator for *C. arabica* (18), *C. canephora* (25), and *C. eugenioides* (24), 7, 3, and 4, respectively, were below the standard threshold (1E-5) for the MAST based motif search used by NLR-Parser. Additionally, 9, 17, and 16 genes present motifs that were detectable using the standard threshold but did not contain at least three consecutive motifs belonging to the NB-ARC domain or presented as motifs in random order. These loci were therefore not annotated in the third step of the NLR-annotator (Supplementary Table 6). After this analysis, we also identified and confirmed genes that were not found by NLR-annotator. Two genes were not found in *C. arabica* (LOC113737176 and LOC113735982), five genes in *C. canephora*, (Cc02_g12220, Cc03_g10360, Cc07_g18800, Cc00_g21910 and Cc00_g35420) and four genes in *C. eugenioides* (LOC113766771, LOC113766774, LOC113766615 and LOC113777141). Supplementary text 1, supplementary table 6 and supplementary figure 1 present additional details from this analysis. After these analyses, it was possible to verify that the NLR annotator showed a sensitivity of 99.8%, 99.4%, and 99.7% for *C. arabica, C. canephora* and *C. eugenioides*, respectively.

As stated above, the overlap analysis also made it possible to detect that the NLR-annotator identified loci that were complete, complete (pseudogenes), partial and partial (pseudogenes) that did not overlap with genes already annotated in reference genomes (Figure 1, Table 1). To further investigate these loci and ensure that they were indeed related to genes encoding NLR proteins, a BLASTx analysis was performed. This analysis showed that of the 298, 240, and 222 loci in *C. arabica, C canephora*, and *C. eugenioides*, only 7, 4, and 6 did not show homology with resistance proteins being found among the five best hits, respectively. (Supplementary Table 7, highlighted in orange).

To describe the sequences that did not show homology to NLRs proteins by BLASTx, a conserved domains analysis was performed (Supplementary Figure 2). Many of these loci do not show homology with NLRs proteins because most of the sequence contains domains related to the family of proteins involved in the activity of transposable elements such as ribonuclease H (RNase H) and reverse transcriptases (RTs). However, it was also possible to identify characteristic domains of NLR proteins such as NB-ARC, Toll / interleukin-1 receptor (TIR), RX-CC_like, and Rx_N, suggesting that these loci cannot be considered false positives. Only three loci did not present characteristic domains, Chr11c_nlr_73_Ca, chr0_nlr_300_Cc and Chr8_nlr_67_Ce and all these loci were partial (pseudogenes). These loci were removed from further analysis. From these results, it was possible to verify that the specificity of the NLR-annotator was 99.9% in all three genomes.

### 3.2 Distribution of NLR loci in the Coffea spp. Genome

Considering all detected loci, in *C. canephora*, chromosomes 3 and 11 have the most significant number of identified loci, including complete, complete (pseudogene), partial and partial (pseudogene). For *C. eugenioides*, chromosomes 3 and 11 also contain many NLR and large numbers of loci were also found on chromosomes 5 and 8. The total number of loci found on chromosome 8 is almost the same as the number identified on 11, however, the number of complete NLR on chromosome 8 is higher, with 16 more loci identified. For *C. arabica*, chromosomes 3 and 11 from the *C. canephora* and *C. eugenioides* subgenomes, respectively, also have the largest number of NLR. This what was also found on chromosome 8 from the *C. eugenioides* subgenoma. For *C. arabica*, the *C. eugenoides* subgenome generally has a slightly higher number of NLRs loci as reported above. The number of loci of this subgenome on chromosomes 8 and 11 stand out in comparison to *C. canephora* subgenome, with 34 and 30 more loci, respectively. The chromosomes with the fewest loci for the three species are 9 and 10, and chromosome 4 specifically for *C. eugenioides*. The number of loci in unmapped sequences (Unassigned) for the *C. canephora* reference genome represent 39.7%, which was much higher than the number of loci found in *C. eugenioides* (12.5%) and *C. arabica* (2,6%) (Figure 2A).

**Figure 2.**
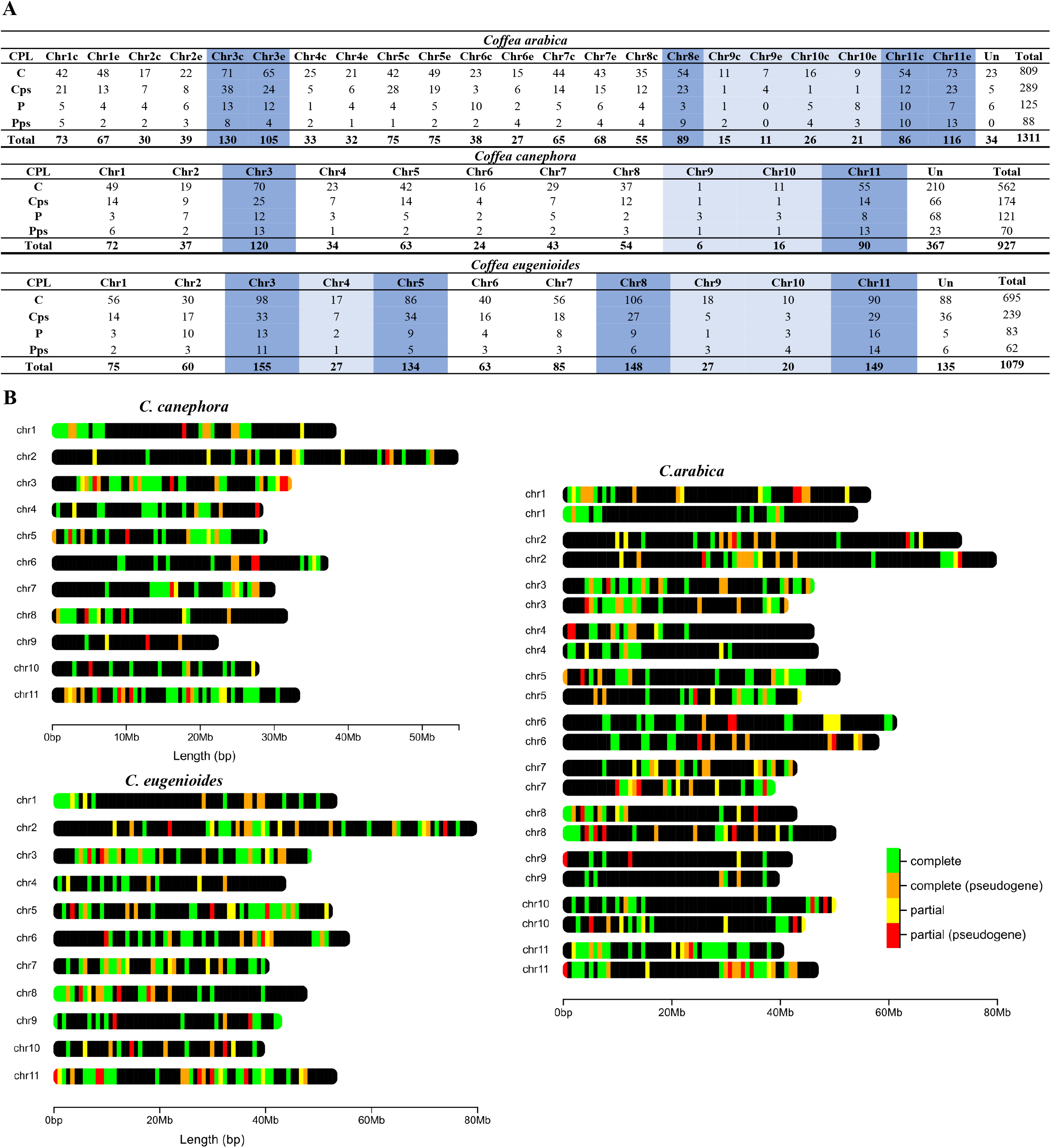
Number and chromosomal distribution of NLR loci in *C. arabica, C. canephora* e *C. eugenioides*. **(A)** The chromosomes with the highest number of NLR loci are highlighted in dark blue, and those with the smallest number of NRL are highlighted in light blue. CPL identifies the completeness of NLR as: C= complete, Cps= complete (pseudogene), P= partial, Pps= partial pseudogene and Un= unassigned. **(B)** The chromosomes represented in *C. arabica* refer to the two subgenomes with the first originating from *C. canephopra* and the second originating from *C. eugenioides*. The chromosomal distribution represented in this figure does not show all loci for each region but identifies all regions that contain NLRs loci. A more detailed view of these chromosomes with the detail of all regions may be found at: https://1drv.ms/u/s!As084N7WlXAIhMZfxevH93zPgU-YhQ?e=0KQ8Iy. Access the link and download the HTML file.

The chromosomal location of these loci in the three species demonstrated that most loci are organized in clusters and are unevenly distributed across the entire chromosome. In addition, there are clusters that have the four different types of loci or at least two types, presenting a stretch of complete, complete (pseudogene), partial and/or partial (pseudogene). Not all loci were clustered, we also found loci of the four types that were physically isolated in chromosomes (Figure 2B).

### 3.3 Gene prediction for complete loci found only by NLR-annotator

Since the NLR-annotator is not a gene predictor but is a tool to annotate loci associated with NLRs, the gene-finding program AUGUSTUS was used to characterize the loci found only by NLR-annotator and that were classified as complete (S3 Table highlighted in orange). This analysis aimed to verify whether these complete loci could be considered potential gene models. This analysis showed that of the 70 and 67 complete loci for *C. arabica* and *C. canephora*, 64 and 66, are potential gene models, respectively. For *C. eugenioides*, all 71 loci were predicted as potential genes. The loci that were not identified as potential genes are in red in supplementary table 3.

### 3.4 Ortholog Groups and Phylogenetic Analysis

From the ortholog group analysis conducted using OrthoMCL, 803 complete loci of *C. arabica*, 561 of *C. canephora* and 695 of *C. eugenioides* were used. Six and 1 loci of *C. arabica* and *C. canephora*, respectively, were removed from analysis because they are complete loci that are not overlapping gene models or were not identified as putative genes by AUGUSTUS analysis. Additionally, 151 tomato loci (out of 326) and 403 potato loci (out of 755) that were classified as complete loci by NLR-annotator as well as 67 reference NLRs and *CED-4* were used. Out of a total of 2681 NLRs, 2038 (76%) were grouped into 593 orthologous groups, hereinafter referred to as orthogroups (Figure 3, Supplementary Table 8). The number of coffee NLRs present within these orthogroups were 591, 427, 584, which represents 73.6%, 76.1% and 84% of the total NLRs found for *C. arabica, C. canephora* and *C. eugenioides*, respectively. Two hundred and seventy-two orthogroups were in single-copies, containing 647 NLRs, of which only 7 are reference NLRs.

**Figure 3.**
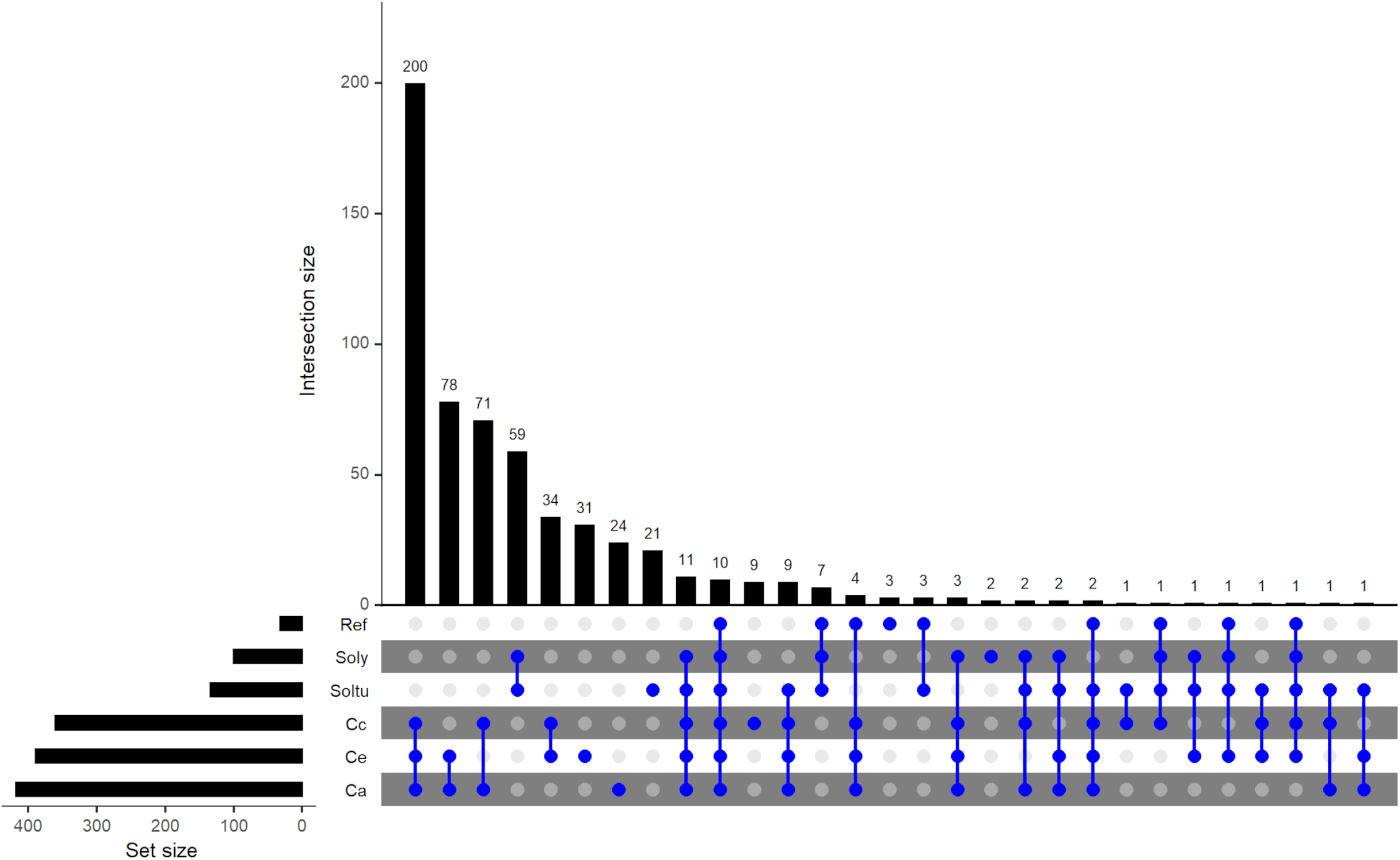
Upset plot of orthologous NLR groups (orthogroups) among five species, *C. arabica* (Ca) *C. canephora* (Cc), *C. eugenioides* (Ce), *S. tuberosum* (Soltu) and *S. lycopersicum* (Soly) and NLRs de referência (Ref). The orthogroups that cluster combinations of species/Ref NLRs is shown by the interconnected blue dots on the bottom panel. Unconnected blue dots show orthogroups that are present within the same species. The ‘Set size’ represents the total number of orthogroups per species/Ref. The ‘intersection size’ shows the number of orthogroups shared between species/Ref or within the same species/Ref. The orthogroups were clustered using OrthoMCL.

There were 10 orthogroups shared by all species and reference NLRs and 11 were shared only among species and not the reference NLRs. As expected, the greatest number of orthogroups were shared among coffee NLRs, 200 orthogroups containing 783 NLRs were shared only among *C. arabica* (296: 163 CaE, 130 CaC e 3 un), *C. canephora* (215) and *C. eugenioides* (272), respectively. The comparison between *C. arabica* NLRs with only one of the diploid species showed that *C. eugenioides* shares a slightly higher number of orthogroups (78) than *C. canephora* (71) and also of NLRs within these orthogroups (orthogroup Ca/Ce = 87/96 NLRs, orthogroup Ca/Cc = 86/74 NLRs). When the comparison was only between the two diploid species, it was observed that 34 orthogroups are shared only between them. The number of orthogroups shared between NLRs of the same coffee species was 31 in *C. eugenioides*, 24 in *C. arabica* and 9 in *C. canephora*.

The number of NLR orthogroups shared only between the three coffee species and one solanum specie was higher among potato (9) than tomato (3) NLRs, however it should be noted that the number of potato NLR in the analysis was almost 3 times larger than the number of tomato NLR. Forty-six orthogroups contain NLRs from at least one coffee species and one solanum species. Fifteen orthogroups were shared between reference NLRs, and at least one coffee and a solanum species, and grouped 21 reference NLRs (Figure 3, Supplementary Table 8, highlighted in light blue). Of these, *C. canephora* and/or *C. eugenioides* are present in three orthogroups with reference genes in which *C. arabica* is absent (ORTHOMCL16: Cc, Soly, Soltu e *Hero;* ORTHOMCL17: Ce, Soly, Soltu e *Rpiblb1;* ORTHOMCL24: Cc, Ce, Soly, Soltu e *VAT*), indicating these orthogroups are not present in the hybrid. Four orthogroups were clustered in only the three coffee species and reference NLRs (ORTHOMCL1, ORTHOMCL19, ORTHOMCL119 and ORTHOMCL199, Figure 3 and Supplementary Table 8, highlighted in dark blue), which contained *Lr10, MLA1, MLA10, MLA13, Mla12, Mla6, Pi36, Pikm2TS, FOM-2, Rdg2a* e *Pm3*. The percentage of orphans (i.e., NLRs not assigned to any ortholog group by OrthoMCL) among coffee NLRs was highest in *C. arabica* (26.4% −212) followed by *C. canephora* (23.9% −134) and *C. eugenioides* (16% −111) (Supplementary Table 9).

The phylogenetic tree of single-copy orthologous NLRs showed that most clades are shared only among coffee species (Figure 4), but it was also possible to observe clades that clustered NLRs from solanum, coffee, and reference. Among the clades that clustered coffee NLRs, 71 presented groupings of orthologs between *C. arabica, C. canephora* and *C. eugenioides*, and most of these are located within the same chromosome. One of these clades, in addition to grouping NLRs of the three coffee species, includes the reference NLR *RPS2* (RESISTANCE to P. SYRINGAE 2) (Figure 4). This clade was supported by a high bootstrap value (100%) and was grouped in the ORTHOMCL45 (Supplementary Table 8). All loci in this cluster were found on chromosome 6 for the three coffee species. Clades that contained NLRs of *C. arabica* and *C. canephora, C. arabica* and *C. eugenioides* and a few *C. canephora* and *C. eugenioides* were also observed. These are within the same chromosome or on different chromosomes.

**Figure 4.**
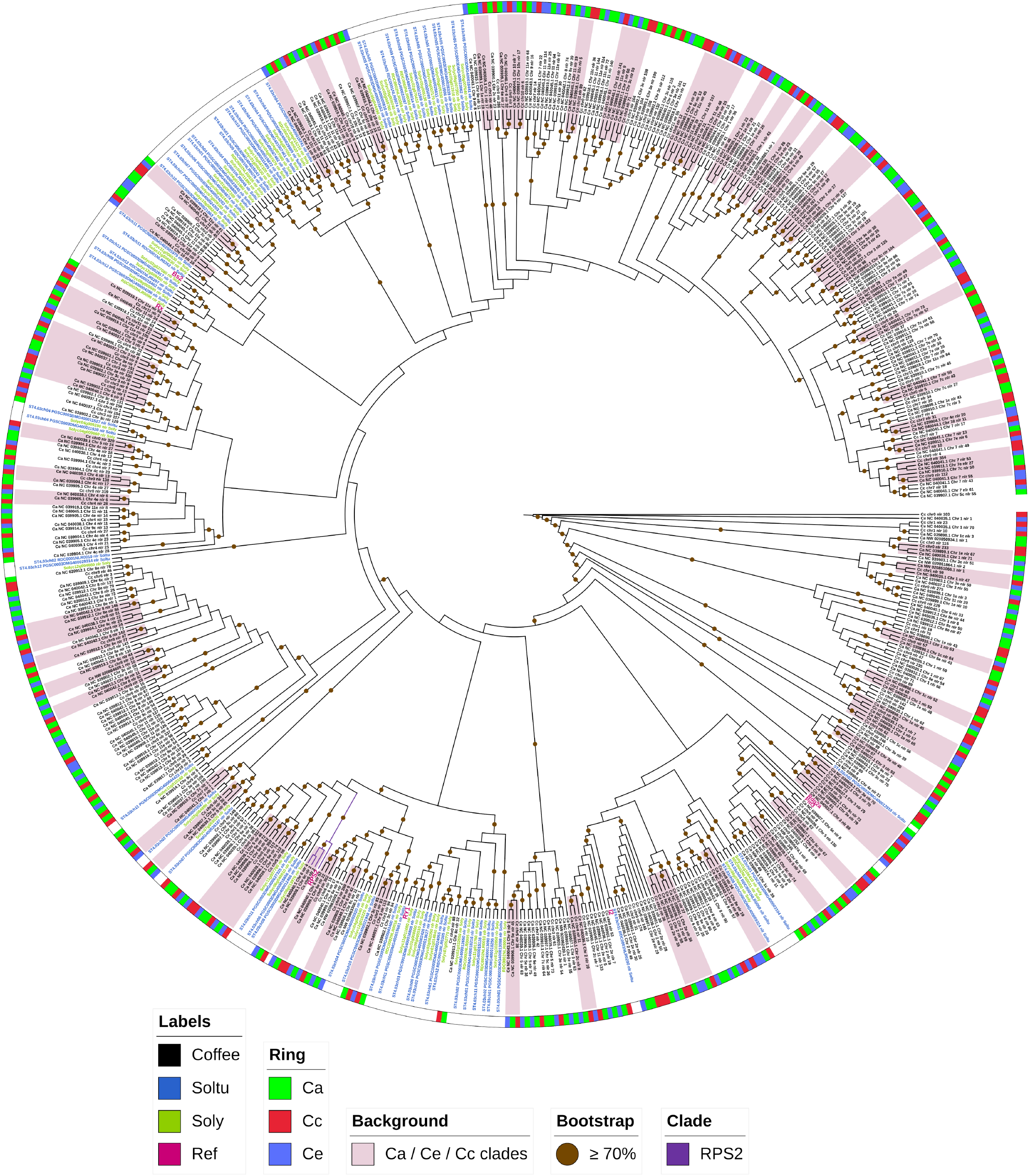
Phylogenetic tree of single-copy NLR orthogroups. The phylogenetic tree was constructed using RAxML and was based on 647 NLRs (domain NB-ARC) that were single-copy orthologs. The colored ring indicates coffee NLR clades, the green color represents *C. arabica* (Ca), red represents *C. canephora* (Cc) and blue represents *C. eugenioides* (Ce). Labels in black are coffee NLRs, green is used for *S. lycopersicum* (Soly), blue is used for *S. tuberosum* (Soltu) and pink indicates reference proteins (Ref). Bootstrap values above 70% are indicated on each branch with a brown circle. The pink background identifies clades that group orthologs of Ca, Cc and Ce. The clade highlighted in purple shows the coffee NLRs and RPS2 grouping.

Phylogenetic analysis for coffee NLRs classification revealed that members of the NLR superfamily are grouped into 2 main groups: TIR-NLR (including TNL and NLs) and non-TNLs (including CNLs and NLs) (Figure 5). NLRs belonging to the non-TNL group outnumbered those in the TNL group in coffee genomes. We also found that the non-TNLs group is divided into 13 subgroups and that all subgroups had NLRs from all studied coffee genomes. The same pattern occurred in the TNL group. Within non-TNLs subgroups it was possible to observe clades with a greater number of NLRs from *C. arabica* that are shared with *C. eugenioides* (bands on the outer ring of the tree with a predominance of green and blue colors). There were exclusive coffee clades as well as clades that contained NLRs that were shared with potato, tomato, and reference NLRs.

**Figure 5.**
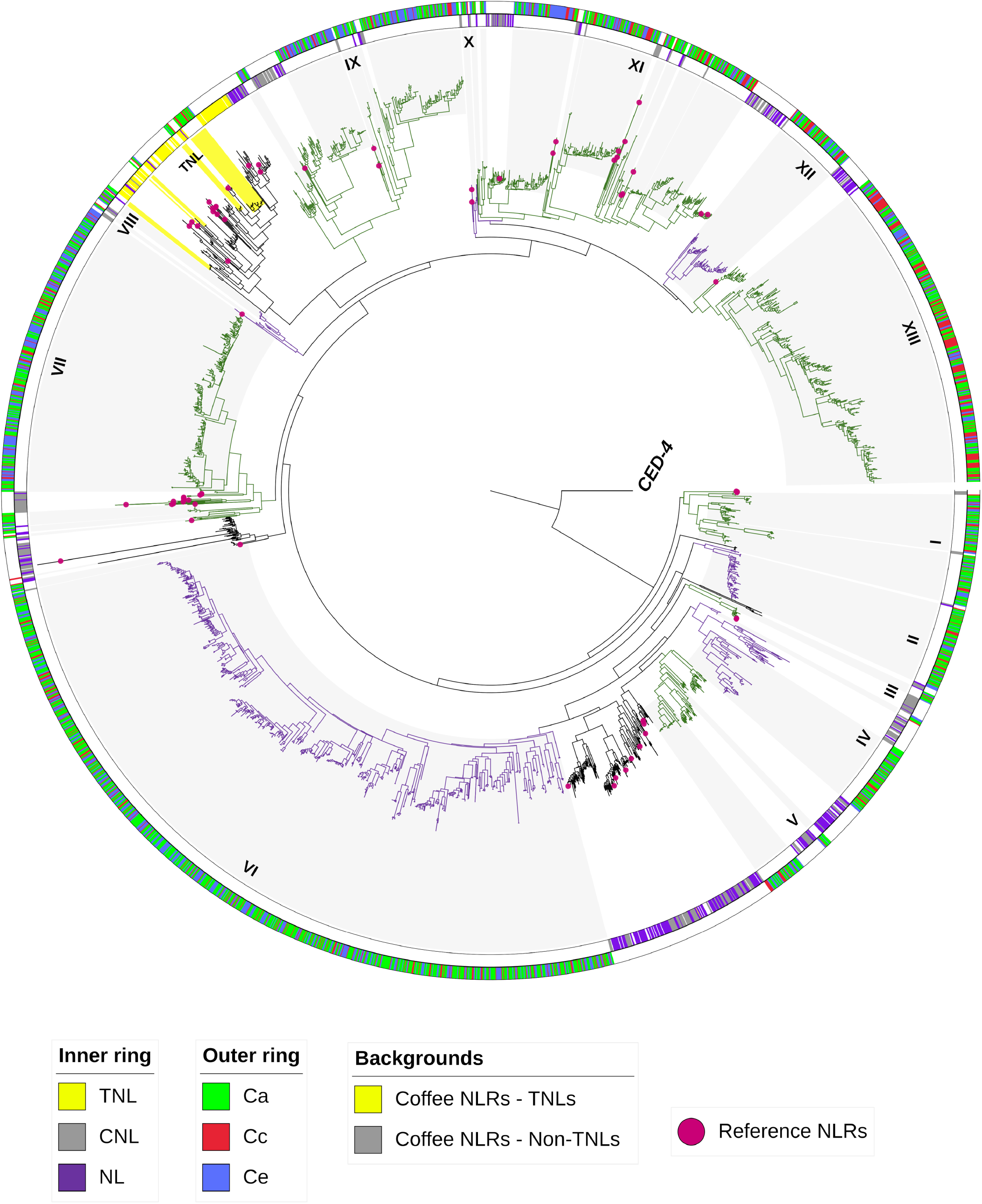
Phylogenetic tree for coffee NLRs classification. NB-ARC domains from 2681 NLRs clustering *C. arabica* (*Ca*), *C. canephora (Cc), C. eugenioides* (Ce), *S. lycopersicum, S. tuberosum*, reference NLRs (pink balls) and CED-4 (root nematode outlier) were used to construct the tree. The tree was constructed using RAxML. The classifications of the reference NLRs and some *S. lycopersicum, S. tuberosum* NLRs were used to classify the coffee NLRs into TNLs and Non TNLs groups (inner ring - TNL = yellow, CNL = gray and NL = Purple). Subgroups in Non TNLs are indicated from I to XIII and alternating colors (green and purple). Gray and yellow background highlight coffee NLRs and the outer ring separates the NLRs for Ca, Cc and Ce in green, red and blue respectively.

## 4 Discussion

### 4.1 NLRs Identification and use of NLR-annotator in coffee genomes

In this study, we investigated loci related to genes of the NLR family in three coffee genomes and compared these loci with NLR from other plants. The annotation of genes in this family is a high priority in plant genome sequencing and annotation projects because losses from pathogens are among the main problems for sustainable agriculture (Steuernagel et al., 2015). A better understanding of disease resistance in crops will provide plant breeders with tools that may be used to produce long-term solutions for dealing with future environmental change. A catalog of NLRs loci, whether complete genes or pseudogenes, within and between species, provides a toolbox for exploring NLRs that has not previously been described (Jones et al., 2016). Given the importance of coffee and the availability of the recent *C. arabica* and *C. eugenioides* genomes, the study of NLRs loci in these species represents an essential source of information for the development of new disease-resistant cultivars.

As NLR-annotator has already been validated in *C. canephora* (Steuernagel et al., 2020), we initially used this genome to ensure the reproducibility of the tool in our study and subsequently applied it with the *C. arabica* and *C. eugenioides* genomes. The tool identified 932 loci NLR for *C. canephora*, which is in accordance with Steuernagel et al., 2020, and also predicted of two distinct NLRs loci within the same genomic position that was previously reported. This repeatable annotation is the result of a complete NB-ARC domain preceded by a truncated NB-ARC domain, which makes the tool use the two NB-ARC domains as distinct seeds to identify two NLRs for the same locus. This has also been reported when this tool was used on the wheat genome (Steuernagel et al., 2020). The sensitivities and specificities of this tool in coffee genomes were extremely high (above 99%). In the *Arabidopsis thaliana* genome, which represents a well-annotated model plant genome, this tool achieved 95% sensitivity, and all loci that were found were validated to be associated with NLRs (specificity of 100%) (Steuernagel et al., 2020). In the Nipponbare reference genome of rice, the detection success rate was 99.2% (Read et al., 2020).

As NLR genes have repeated and clustered genomic distributions in plants, their representation in genomes using standard gene callers can be underestimated (Jupe et al., 2013; Steuernagel et al., 2015, 2020). In addition to the high rate of specificity and sensitivity, the NLR-annotator allowed for the identification of complete loci for coffee in regions distinct from the gene models already annotated in the reference genomes. This study, therefore, increased the number of putative NLR genes detected in the reference genomes of coffee species. The complete loci identified by NLR-annotator that did not overlap the reference genome annotation have also been reported in rice (Read et al., 2020). It is also relevant to highlight that as this tool is not limited to searching for functional genes, the complete (pseudogenes) loci that did not overlap with annotations of the reference genome were also identified for *C. arabica* (90), *C. canephora* (56), and *C eugenioides* (65). The location of these loci also represent an important resource, since non-functional alleles identified in sequenced accessions may represent functional NLRs in other individuals of the same species (Steuernagel et al., 2020). The caturra cultivar (*C. arabica*) sequenced and used in this study, for example, is used as a susceptible control to differentiate *Hemileia vastatrix* races among differential clones (Zambolim and Caixeta, 2021). Pseudogene regions in this genome may indicate functional genes present in other coffee cultivars.

Our data showed that 18 of the 20 loci found only by NLR-annotator, that did not present homology to NLRs proteins by BLASTx analysis, have protein domains involved in the activity of transposable elements (TE). It is known that TE are abundant in plant genomes and that they play an important role in adaptive evolution and contribute to the evolutionary dynamics of plant-pathogen interactions (Malacarne et al., 2012; Zhang et al., 2014; Kim et al., 2017). Many R genes are flanked by TE, which in addition to being sources of genetic variability, are involved in suppressing or increasing the expression of these genes (Seidl and Thomma, 2017). The Ty3-gypsy-like TE performance has been reported in a region around the *S_H_3* locus associated with CLR resistance. This TE has been described in *C. arabica* subgenomes, replacing the orthologous counterpart of *C. canephora* with that of *C. eugenioides* (homoeologous non-reciprocal transposition - HNRT) (Cenci et al., 2012). Moreover, there is evidence of functional R genes that have evolved through TE-mediated duplications (Seidl and Thomma, 2017), which demonstrates their importance in the evolutionary changes and expansion of NLR receptors and justifies the presence of domains related to TE in the studied loci (Zhang et al., 2014; Kim et al., 2017).

### 4.2 Distribution of NLR loci in the *Coffea spp*. Genome

Although *C. arabica* results from a natural interspecific hybridization event between *C. canephora* and *C. eugenioides*, the number of loci found was not proportional to the sum of the two subgenomes, showing that the hybrid has a relatively smaller number of NLRs loci. The *C. canephora* genome size is about 690 Mbp, and the *C. eugenioides* is 665 Mbp (Noirot et al., 2003; Clarindo and Carvalho, 2011; Hamon et al., 2015). The *C. arabica* genome, on the other hand, is slightly smaller than the sum of its two combined parental genomes (about 1276 Mbp) (Hamon et al., 2015). This may explain the smaller number of NLRs in this species. Genome contraction is common in amphidiploids, which may be related to chromosomal rearrangements, including duplication, insertions, and deletions after initial hybridization (Hamon et al., 2015). An example of the number of NLRs being smaller than the sum of the corresponding parents was reported in *Brassica juncea* (Indian mustard), a species formed by hybridization between the diploid Brassica species of *B. rapa*, and *B. nigra* (Inturrisi et al., 2020). Moreover, differences in the genome assembly quality may also have interfered with the identification of NLR loci.

Among the three species analyzed, the only one with a genome-wide NLR study already reported is *C. canephora* (Denoeud et al., 2014). The NLR gene data from the previous study agrees with much of our findings. A large number of NLR loci in unanchored scaffolds for this species has also been described. Here 210 complete NLR loci were identified in unanchored scaffolds for *C. canephora*, while in the first description of the manually curated genes, 213 were not mapped. The number of mapped NLR genes was 348, while in our study, there were 352 complete loci. In *C. canephora*, it has also been reported that NLRs genes are located on all chromosomes, but with an increased number found on chromosomes 1, 3, 5, 8, 11, which together represented 70.1% of the mapped NLR genes. Although we have highlighted chromosomes 3 and 11 as having a greater number of NLR, chromosomes 1, 5, and 8 also contain large numbers of NLR loci in the three species studied here. Together, all these chromosomes represent 68.2, 71.4 and 70.0% of the total of NLR loci mapped for *C. arabica, C. canephora*, and *C. eugenioides*, respectively. Moreover, the low number of NLR genes on chromosomes 9 and 10 had been previously reported was confirmed in this study (Denoeud et al., 2014). These comparisons show that the three species display a conserved pattern with regards to the chromosomal distribution of NLR loci.

The NLR loci found in the three studied coffee species are arranged in clusters that group complete loci, pseudogenes and partial. These genes tend to group together and provide birth-and-death events for functional NLRs (Ling et al., 2021). In these clusters it is possible to find tandem gene duplications, recombination hotspots or active transposon elements functioning as a reservoir of genetic variation to generate specificity for new pathogen variants (Michelmore and Meyers, 1998; Zhang et al., 2014). Within plant genomes many R genes have been found to reside in clusters (Jupe et al., 2012; Andolfo et al., 2014, 2021; Seo et al., 2016; Zheng et al., 2016; Read et al., 2020). The *S_H_3* locus in coffee, for example, corresponds to a complex cluster of multiple genes, including CNL-like NLR genes (Ribas et al., 2011; Cenci et al., 2012). The number of complete or functional loci in plants represents a fraction of the total number of loci found (Jupe et al., 2012; Seo et al., 2016). This happens precisely because the evolutionary dynamics within these clusters favor the coexistence of functional genes, pseudogenes, and partial genes, which differ between plants in consequence of evolutionary routes for certain pathosystems.

Recent discoveries show that NLRs can be multi domain receptors, that is they present domains integrated to the canonical form NLR or TNL/CNL (Bailey et al., 2018; Wang et al., 2021). Knowing regions of the genome that have this canonical form can facilitate the description of non-canonical integrated domains that are upstream or downstream from the more conserved region (Monteiro and Nishimura, 2018). Other relevant information is that activation of NLRs often happens in complexes and there is evidence that truncated NLRs can form heterocomplexes with complete NLRs, or may act as the main receptors in defense activation (Monteiro and Nishimura, 2018). NLRs truncated as *CbCN* (*Capsicum baccatum* – CC-NB-ARC), and *TN2* (TIR-NB-ARC) act in resistance to pathogens (Zhao et al., 2015; Son et al., 2021). This evidence reinforces the importance of knowing loci related to R genes, whether complete, pseudogenes or partial.

### 4.3 Ortholog groups and Phylogenetic Analyses

The genus *Coffea* belongs to the Rubiaceae family, which is in the asterid clade that also contains the Solanaceae family. Many studies have used species of the genus *Solanum* to obtain insights into the genomic and evolutionary architecture of coffee (Lin et al., 2005; Lefebvre-Pautigny et al., 2010; Denoeud et al., 2014). Species of the genus *Solanum* have also been used as models for understanding the molecular processes related to plant-pathogen interaction. This supports their use for comparative approaches to lead to discoveries of NLR loci or functionally important gene families in coffee (Andolfo et al., 2021). Our results showed that of the 17 reference NLRs that have been cloned and characterized in species of the genus *Solanum* and that were also used in this study, 8 of them are present in shared orthogroups with coffee (Supplementary Table 2 and Supplementary Table 8), being 1 TNL (*Gro1-4*) and 7 CNL (*Hero, Prf, Rpi-blb1, Rx2, Sw-5, Tm-2a, Tm-2*). These genes have been found to be involved in resistance to a diverse group of pathogens including viruses, oomycetes, bacteria and nematodes (Bendahmane et al., 2000; Van Der Vossen et al., 2003; Paal et al., 2004; Andolfo et al., 2021). In total, 46 orthogroups were shared between coffee and solanum indicating that these orthologs were probably present before the speciation of these two groups. Reference NLRs characterized in species such as *Hordeum vulgare, Oryza sativa, Triticum aestivum, Glycine max, Arabidopsis thaliana* e *Cucumis melo* also share othogroups with coffee NLRs and all of these NLR belong to the CNL class (Supplementary Table 2 and Supplementary Table 8). These orthogroups are important as they indicate roles that may be inferred and further investigated in coffee. An interesting orthogroup that obtained high support in the phylogenetic tree was the one that clustered the *RPS2* reference NLR as well as NLRs present in all three coffee species. *RPS2 is* a resistance gene of *Arabidopsis thaliana* that confers resistance to *Pseudomonas syringae* bacteria that express the *avrRpt2* avirulence gene (Bent et al., 1994; Mindrinos et al., 1994).

In general, plant species exhibit differences in the number of NLR genes contained within their genomes. Amplification of certain groups has been detected (Seo et al., 2016). This diversity has not been associated with genome size or phylogenetic relationships, but is related to the specialization of each particular host (Wan et al., 2013; Lozano et al., 2015; Seo et al., 2016; Zheng et al., 2016; Steuernagel et al., 2020). In all three coffee species, a set of orphan genes and orthogroups that share NLRs within the same species were detected. In tomato, 45 of ~320 NLRs sequences are more similar to each other than to any other sequences compared (Andolfo et al., 2021) and orthogroups that share NLRs within the same species of *Solanum* are attributed to duplication events that generate different gene repertoires and result in species-specific subfamilies (Seo et al., 2016).

The single-copy ortrogroups provide more reliable results for interpreting evolutionary processes between groups of evaluated genes by allowing for the identification of true orthologs between different groups of plants (Zimmer et al., 2007; Duarte et al., 2010). The results from the single-copy ortrogroups tree showed that some orthologous NLRs seem to have a common ancestor only among coffee species. The *S_H_3* locus, for example, was described as being shared only among coffee species suggesting that the ancestral copy *S_H_3*-CNL was inserted into the *S_H_3* locus after the divergence of the *Solanum* and *Coffea* lineages (Ribas et al., 2011). The clades that clustered orthologous NLRs from *C. arabica, C. canephora* and *C. eugenioides* probably represent NLR present in both ancestral diploids genomes, which were passed to *C. arabica* genome. On the other hand, clades that clustered only *C. canephora* and *C. eugenioides* loci may represent ancestral NLRs that were lost in *C. arabica* or that underwent so many modifications in this species that makes it difficult to find homology between these NLRs. These NLR may provide valuable resistance mechanisms that are not present in the *C. arabica* hybrid. Nucleotide level changes, such as deletions, insertions and rearrangements have been observed in coffee RGA (Resistance gene analogues) (Noir et al., 2001; Hendre et al., 2011). It is also known that it is very likely that the sequenced genotypes of *C. canephora* and *C. eugenioides* present significant differences from the ancestral donors of *C. arabica* subgenomes, which may explain the lack of homology in certain NLR groups (Cenci et al., 2012).

The NB-ARC is the most conserved domain in the NLR gene family. Despite the conservation of this domain, it is possible to distinguish the TIR (TNL) and non-TIR classes based on different residues inside the motifs present in this region (Jones et al., 2016; Shao et al., 2019; Van Ghelder et al., 2019). Therefore, this domain has been used to describe the phylogenetic relationships between the sequences of this group and classify them (Andolfo et al., 2014). The classification of NLRs in coffee revealed that the non-TNL class were present in greater numbers than those of the TNL group in each of the three analyzed coffee genomes. It is known that non-TNL genes that include many CNL are widely distributed in monocots and dicots, while TNL are mainly found in dicots (McHale et al., 2006; Zheng et al., 2016). The low frequency of TNLs in coffee agrees with results found for species of the solanum group, such as pepper, tomato and potato (Andolfo et al., 2014; Seo et al., 2016). Our results are also consistent with the low frequency of TNLs found in previous studies of coffee (Hendre et al., 2011; Denoeud et al., 2014). Thus, it is possible to suggest that, as in solanum, non-TNLs represent an important repertoire of resistance genes in coffee. Additionally, the TNL group and the non-TNLs subgroups contained NLRs from *C.arabica, C. canephora* and *C. eugenioides*, indicating conservation of the NLR classes across coffee genomes and suggesting that all subgroups were present in a common ancestor, similar to what has been described for comparisons of species within the solanum group (Seo et al., 2016).

In the two phylogenetic trees analyzed, the clades group coffee NLRs that are mostly present in the same chromosomes but groupings of NLRs present on different chromosomes were also detected. Genes located on the same chromosome tend to group into subclades in the phylogenetic tree. However, rearrangement events of the chromosomes can affect NLR loci and modify their genomic order or location (Cenci et al., 2012; Zheng et al., 2016; Ling et al., 2021).

Considering the relevance of coffee, few studies have been conducted addressing the identification of NLR in genomes of this crop. RGA studies using degenerate primers for NB-ARC region have already been performed (Noir et al., 2001; Hendre et al., 2011; Kumar, 2012), in addition to studies in *S_H_3* locus (Cenci et al., 2010, 2012; Lashermes et al., 2010), but very little is known about the NLR family in cultivated (*C.arabica* e *C. canephora*) and uncultivated coffee species (such as *C. eugenioides*). This is the first study focused on genome wide identification of NLRs in the *C. arabica* genome, and also adds information to the existing report for the *C. canephora* genome (Denoeud et al., 2014). The Genome-wide identification of coffee NLRs allows for more in-depth molecular studies provides further information for identifying candidate genes for cloning and subsequent functional validation of NLR genes while also expanding the range of NLR that are available for breeding of this crop (Seo et al., 2016).

## 5 Conclusion

Our analysis showed that the use of a specific pipeline for resistance genes was efficient in detecting NLR loci in the studied coffee genomes and increased the information available for the location of these loci in *C. arabica, C. canephora* and *C. eugenioides*. The NLR loci in the three coffee species are unevenly distributed across all chromosomes and are mostly arranged in clusters. The number of loci in *C. arabica* is less than the sum of the NLR loci from the parents of this hybrid. Single-copy NLR orthogroups investigated in a phylogenetic tree identified orthologous NLRs that are shared between *C. arabica* and the parental genomes as well as NLR that were possibly lost. Coffee NLRs and some functionally characterized NLRs share common orthogroups, which provides clues to the functionality of coffee NLRs, and paves the way for further investigation. Loci from the NLR family are subdivided into two main groups in coffee: TIR-NLR (TNL) and non-TNL, but non-TNLs are present in greater number and seem to represent an important repertoire of resistance genes in coffee.

## 7 Author Contributions

MLi, GS, and MF: conceptualization. MLi, MLu, GS, JC, MF, and JT: methodology. MLi, JC: software, formal analysis, investigation. MLi writing – original draft preparation. MLi, MLu, GS, JC, MF, and JT: visualization. MLi, MLu, GS, JC, MF, and JT: validation. MLu, and MF: resources, supervision, project administration, and funding acquisition. MLi, MLu, GS, JC, MF, and JT: data curation; writing – review and editing. All authors have read and agreed to the published version of the manuscript.

## 9 Acknowledgments

We are grateful to the Universidade Federal de Lavras, the Universidade de Brasília, the University of Florida, U.S. Department of Agriculture’s Agriculture and Food Research Initiative grant FLA-PLP006039, University of Florida’s Institute for Food and Agricultural Science, and Corn Breeding and Genomics program for supporting this collaborative research project.

