## Supplementary Figure 1 for "Genome-wide identification, characterization, and comparative analysis of NLR resistance genes in *Coffea spp*"

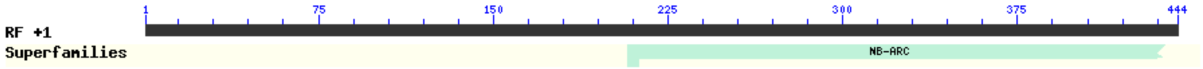


| ID | Name | Accession | Description | Interval | E-value |
|---|---|---|---|---|---|
| **LOC113737176_Ca** | NB-ARC super family | cl26397 | NB-ARC domain | 208-438 | 8.92e-11 |

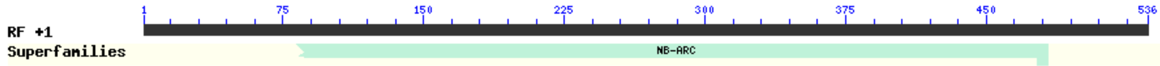


| ID | Name | Accession | Description | Interval | E-value |
|---|---|---|---|---|---|
| **LOC113735982_Ca** | NB-ARC super family | cl26397 | NB-ARC domain | 82-483 | 1.90e-11 |

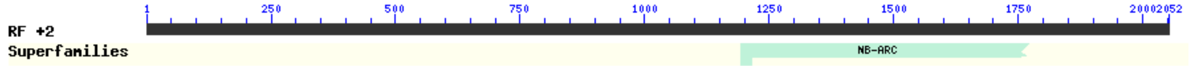


| ID | Name | Accession | Description | Interval | E-value |
|---|---|---|---|---|---|
| **Cc02_g12220_Cc** | NB-ARC super family | cl26397 | NB-ARC domain | 1193-1771 | 3.08e-08 |

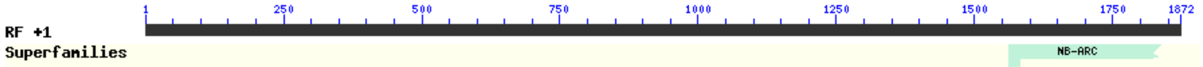


| ID | Name | Accession | Description | Interval | E-value |
|---|---|---|---|---|---|
| **Cc03_g10360_Cc** | NB-ARC super family | cl26397 | NB-ARC domain | 1561-1836 | 7.70e-13 |

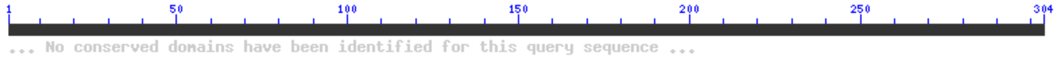


| ID | Name | Accession | Description | Interval | E-value |
|---|---|---|---|---|---|
| **Cc07_g18800_Cc** | - | - | - | - | - |

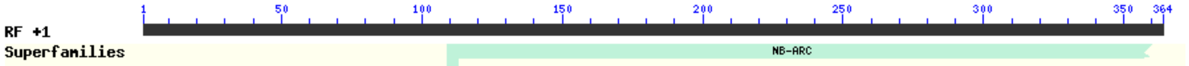


| ID | Name | Accession | Description | Interval | E-value |
|---|---|---|---|---|---|
| **Cc00_g21910_Cc** | NB-ARC super family | cl26397 | NB-ARC domain | 109-360 | 7.62e-08 |

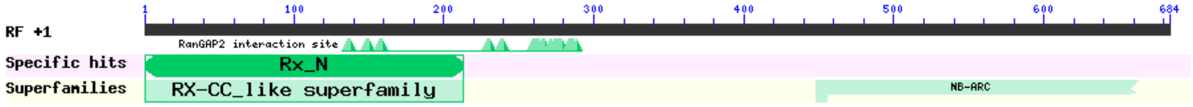


| ID | Name | Accession | Description | Interval | E-value |
|---|---|---|---|---|---|
| **Cc00_g35420_Cc** | Rx_N | pfam18052 | Rx N-terminal domain | 1-213 | 1.29e-17 |
| | NB-ARC super family | cl26397 | NB-ARC domain | 448-663 | 1.78e-10 |

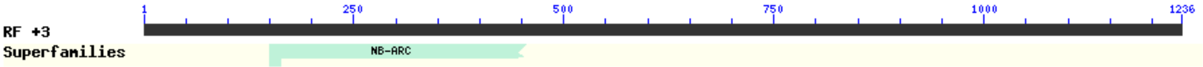


| ID | Name | Accession | Description | Interval | E-value |
|---|---|---|---|---|---|
| **LOC113766615_Ce** | NB-ARC super family | cl26397 | NB-ARC domain | 150-455 | 1.53e-11 |

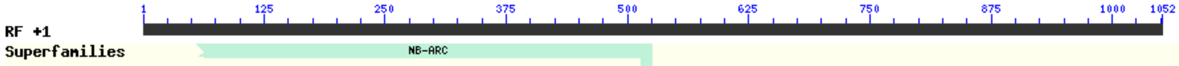


| ID | Name | Accession | Description | Interval | E-value |
|---|---|---|---|---|---|
| **LOC113766771_Ce** | NB-ARC super family | cl26397 | NB-ARC domain | 55-525 | 9.82e-07 |

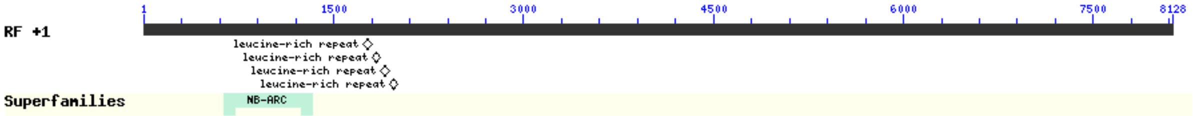


| ID | Name | Accession | Description | Interval | E-value |
|---|---|---|---|---|---|
| **LOC113766774_Ce** | NB-ARC super family | cl26397 | NB-ARC domain | 637-1326 | 9.69e-11 |

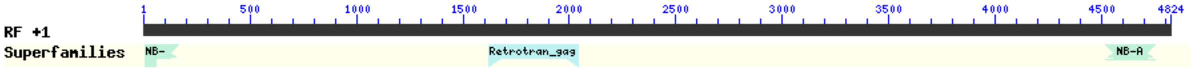


| ID | Name | Accession | Description | Interval | E-value |
|---|---|---|---|---|---|
| **LOC113777141_Ce** | Retrotran_gag_2 super family | cl26047 | gag-polypeptide of LTR copia-type; This family is found in Plants and fungi | 1624-2040 | 2.22e-14 |
| | NB-ARC super family | cl26397 | NB-ARC domain | 7-165 | 1.58e-10 |
| | NB-ARC super family | cl26397 | NB-ARC domain | 4519-4749 | 7.73e-10 |

**Supplementary Figure 1**. **Conserved domains analysis for genes not found by the NLR-annotator in the coffee genomes**. For the analysis, nucleotide sequences were used. The table shows the list of domain hits found for each of the sequences. The conserved domains were detected using NCBI Conserved Domain Database and the graphical summary was set to concise view. If the alignment omitted more than 20% of the either the N- or C-terminus or both, the partial nature of the hit is indicated in the graphical display as domain with jagged edges. At the end of each ID, the source genome is identified, Ca: *C. arabica*, Cc: *C. eugenioides* and Ce: *C. eugenioides*. RF: reading frame.
