## Supplementary Figure 2 for "Genome-wide identification, characterization, and comparative analysis of NLR resistance genes in *Coffea spp*"

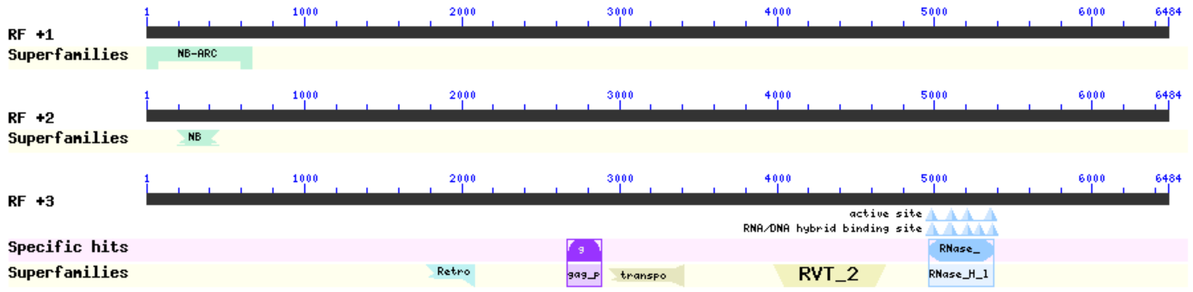


| Chr_4c_nlr_22_Ca partial | | | | |
|---|---|---|---|---|
| **Name** | **Accession** | **Description** | **Interval** | **E-value** |
| NB-ARC super family | cl26397 | NB-ARC domain | 1-663 | 1.67e-06 |
| NB-ARC super family | cl26397 | NB-ARC domain | 194-457 | 4.21e-12 |
| RNase_HI_RT_Ty1 | cd09272 | Ty1/Copia family of RNase HI in long-term repeat retroelements | 4965-5378 | 2.30e-60 |
| RVT_2 super family | cl06662 | Reverse transcriptase (RNA-dependent DNA polymerase) | 3978-4682 | 8.87e-51 |
| Retrotran_gag_2 super family | cl26047 | gag-polypeptide of LTR copia-type; This family is found in Plants and fungi | 1779-2075 | 5.08e-16 |
| transpos_IS481 super family | cl41329 | IS481 family transposase | 2934-3404 | 3.45e-12 |
| gag_pre-integrs | pfam13976 | GAG-pre-integrase domain; This domain is found associated with retroviral insertion elements | 2670-2885 | 3.14e-11 |

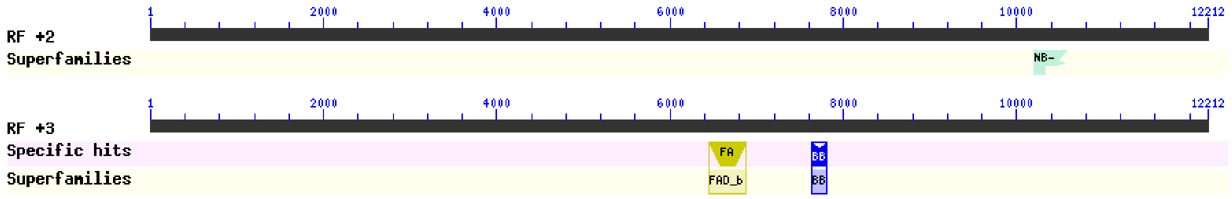


| Chr_3c_nlr_44_Ca complete (pseudogene) | | | | |
|---|---|---|---|---|
| **Name** | **Accession** | **Description** | **Interval** | **E-value** |
| NB-ARC super family | cl26397 | NB-ARC domain | 10199-10585 | 2.48e-10 |
| FAD_binding_4 | pfam01565 | FAD binding domain; This family consists of various enzymes that use FAD as a co-factor | 6453-6872 | 5.37e-19 |
| BBE | pfam08031 | Berberine and berberine like; This domain is found in the berberine bridge and berberine | 7632-7805 | 5.77e-14 |

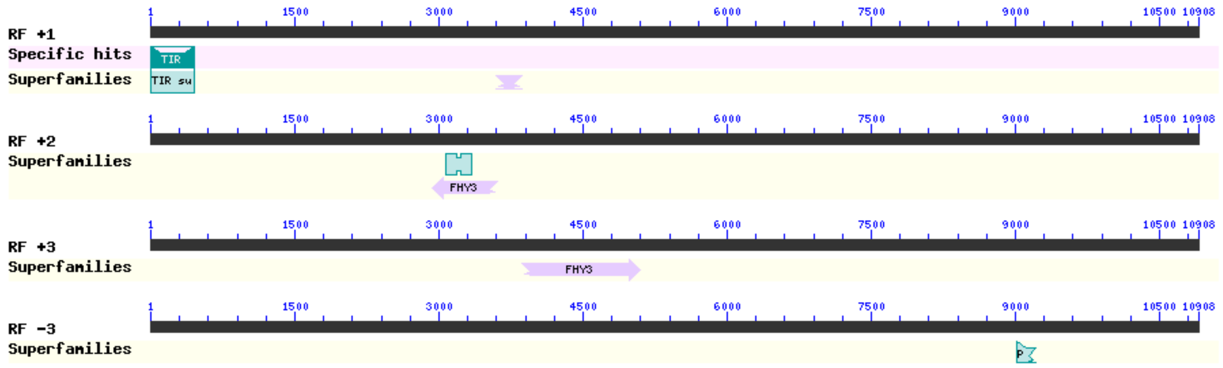


| Chr_10c_nlr_18_Ca partial (pseudogene) | | | | |
|---|---|---|---|---|
| **Name** | **Accession** | **Description** | **Interval** | **E-value** |
| TIR | pfam01582 | TIR domain; The Toll/interleukin-1 receptor (TIR) | 4-450 | 1.05e-54 |
| FHY3 super family | cl31971 | Protein FAR-RED ELONGATED HYPOCOTYL 3 | 3598-3861 | 5.24e-22 |
| FAR1 super family | cl40636 | FAR1 DNA-binding domain | 3074-3334 | 1.24e-19 |
| FHY3 super family | cl31971 | Protein FAR-RED ELONGATED HYPOCOTYL 3 | 2936-3610 | 7.11e-17 |
| FHY3 super family | cl31971 | Protein FAR-RED ELONGATED HYPOCOTYL 3 | 3861-5096 | 1.62e-62 |
| PH-like super family | cl17171 | Pleckstrin homology-like domain; The PH-like family includes the PH domain | 9019-9207 | 4.83e-08 |

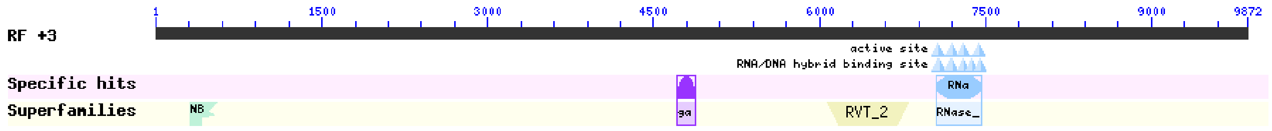


| Chr_8c_nlr_13_Ca complete (pseudogene) | | | | |
|---|---|---|---|---|
| **Name** | **Accession** | **Description** | **Interval** | **E-value** |
| RVT_2 super family | cl06662 | Reverse transcriptase (RNA-dependent DNA polymerase) | 6069-6800 | 2.16e-76 |
| RNase_HI_RT_Ty1 | cd09272 | Ty1/Copia family of RNase HI in long-term repeat retroelements; Ribonuclease H (RNase H) | 7053-7469 | 3.08e-74 |
| gag_pre-integrs | pfam13976 | GAG-pre-integrase domain; This domain is found associated with retroviral insertion elements | 4710-4874 | 1.19e-08 |
| NB-ARC super family | cl26397 | NB-ARC domain | 309-554 | 2.54e-06 |

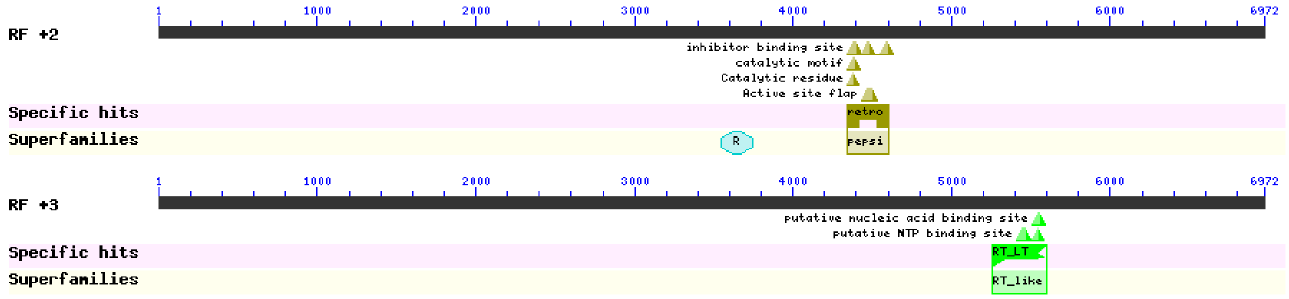


| Chr_11c_nlr_73_Ca partial (pseudogene) | | | | |
|---|---|---|---|---|
| Name | Accession | Description | Interval | E-value |
| retropepsin_like | cd00303 | Retropepsins; pepsin-like aspartate proteases | 4346-4606 | 5.67e-11 |
| Retrotrans_gag super family | cl29674 | Retrotransposon gag protein; Gag or Capsid-like proteins from LTR retrotransposons. | 3542-3745 | 1.56e-07 |
| RT_LTR | cd01647 | RT_LTR: Reverse transcriptases (RTs) | 5256-5597 | 2.09e-40 |

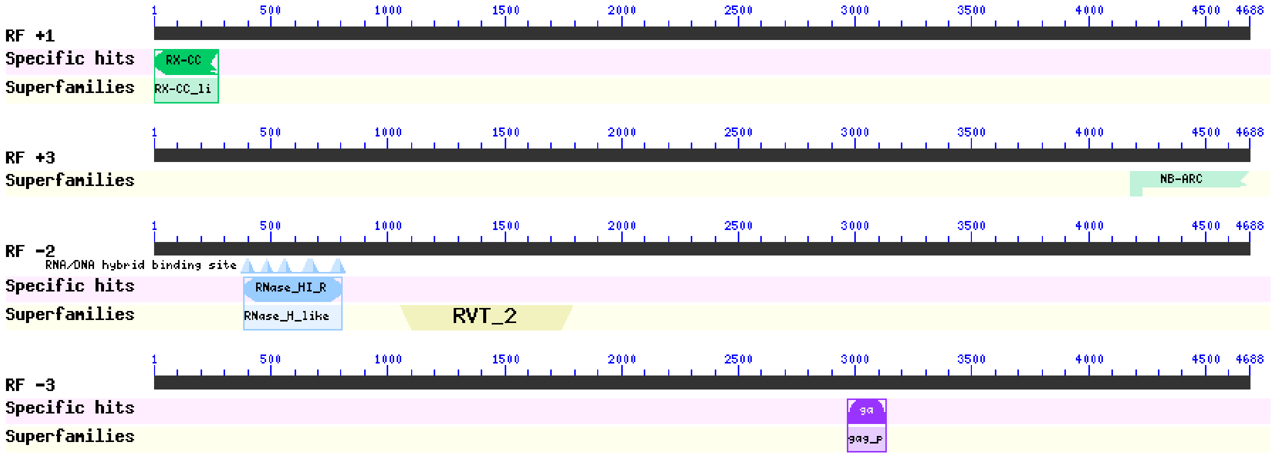


| Chr_6e_nlr_10_Ca partial (pseudogene) | | | | |
|---|---|---|---|---|
| Name | Accession | Description | Interval | E-value |
| RX-CC_like | cd14798 | Coiled-coil domain of the potato virux X resistance protein and similar proteins | 1-273 | 3.80e-15 |
| NB-ARC super family | cl26397 | NB-ARC domain | 4176-4685 | 2.49e-12 |
| RNase_HI_RT_Ty1 | cd09272 | Ty1/Copia family of RNase HI in long-term repeat retroelements; Ribonuclease H (RNase H) | 386-802 | 4.93e-78 |
| RVT_2 super family | cl06662 | Reverse transcriptase (RNA-dependent DNA polymerase) | 1055-1786 | 1.73e-68 |
| gag_pre-integrs | pfam13976 | GAG-pre-integrase domain; This domain is found associated with retroviral insertion elements | 2965-3132 | 1.08e-06 |

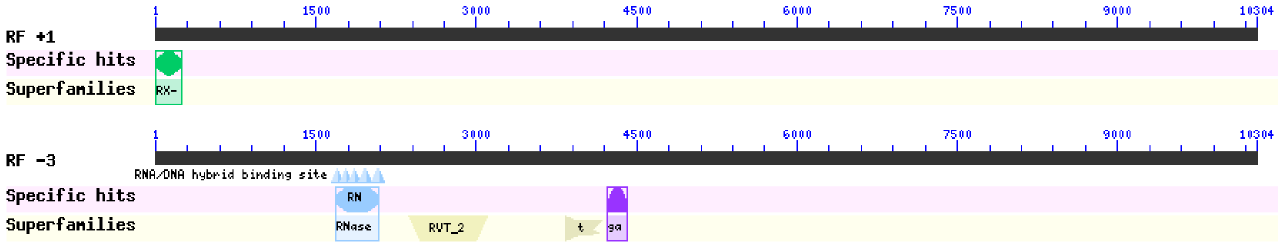


| Chr_1e_nlr_61_Ca complete (pseudogene) | | | | |
|---|---|---|---|---|
| Name | Accession | Description | Interval | E-value |
| Rx_N | pfam18052 | This entry represents the N-terminal domain found in many plant resistance proteins | 1-243 | 9.30e-08 |
| RVT_2 super family | cl06662 | Reverse transcriptase (RNA-dependent DNA polymerase) | 2365-3102 | 1.26e-70 |
| RNase_HI_RT_Ty1 | cd09272 | Ty1/Copia family of RNase HI in long-term repeat retroelements; Ribonuclease H (RNase H) | 1678-2091 | 3.28e-64 |
| transpos_IS481 super family | cl41329 | IS481 family transposase | 3832-4170 | 7.73e-17 |
| gag_pre-integrs | pfam13976 | GAG-pre-integrase domain; This domain is found associated with retroviral insertion elements | 4222-4410 | 1.42e-12 |

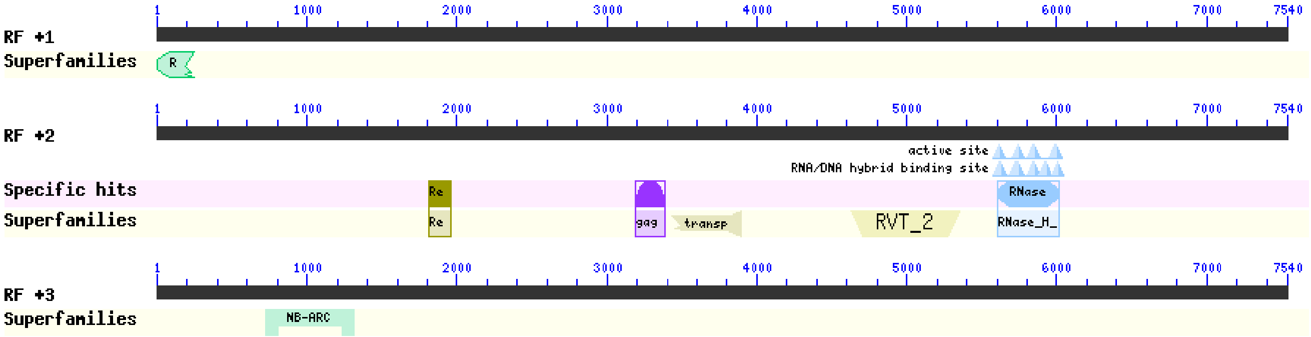


| Chr_11e_nlr_43_Ca complete (pseudogene) | | | | |
|---|---|---|---|---|
| Name | Accession | Description | Interval | E-value |
| RX-CC_like super family | cl36576 | Coiled-coil domain of the potato virus X resistance protein and similar proteins | 1-249 | 9.15e-09 |

| | | | | |
|---|---|---|---|---|
| RVT_2 super family | cl06662 | Reverse transcriptase (RNA-dependent DNA polymerase) | 4622-5350 | 3.81e-78 |
| RNase_HI_RT_Ty1 | cd09272 | Ty1/Copia family of RNase HI in long-term repeat retroelements; Ribonuclease H (RNase H) | 5600-6019 | 4.36e-77 |
| Retrotran_gag_3 | pfam14244 | gag-polypeptide of LTR copia-type; This family is found in Plants and fungi | 1814-1954 | 4.42e-15 |
| transpos_IS481 super family | cl41329 | IS481 family transposase | 3431-3889 | 2.46e-10 |
| gag_pre-integrs | pfam13976 | GAG-pre-integrase domain; This domain is found associated with retroviral insertion elements | 3191-3382 | 1.51e-08 |
| NB-ARC super family | cl26397 | NB-ARC domain | 723-1313 | 1.69e-39 |

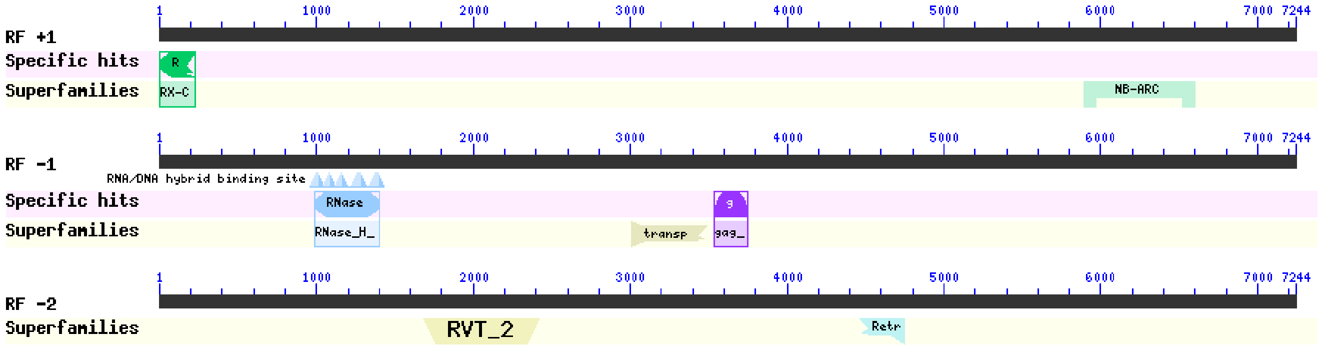


| Chr_10e_nlr_2_Ca complete (pseudogene) | | | | |
|---|---|---|---|---|
| **Name** | **Accession** | **Description** | **Interval** | **E-value** |
| NB-ARC super family | cl26397 | NB-ARC domain | 5896-6600 | 5.29e-44 |
| RX-CC_like | cd14798 | Coiled-coil domain of the potato virus X resistance protein and similar proteins | 1-225 | 1.03e-16 |
| RNase_HI_RT_Ty1 | cd09272 | Ty1/Copia family of RNase HI in long-term repeat retroelements; Ribonuclease H (RNase H) | 993-1400 | 8.74e-66 |
| gag_pre-integrs | pfam13976 | GAG-pre-integrase domain; This domain is found associated with retroviral insertion elements | 3534-3746 | 6.29e-12 |
| transpos_IS481 super family | cl41329 | IS481 family transposase | 3012-3482 | 7.08e-11 |
| RVT_2 super family | cl06662 | Reverse transcriptase (RNA-dependent DNA polymerase) | 1679-2419 | 1.18e-48 |
| Retrotran_gag_2 super family | cl26047 | gag-polypeptide of LTR copia-type; This family is found in Plants and fungi | 4460-4741 | 2.61e-09 |

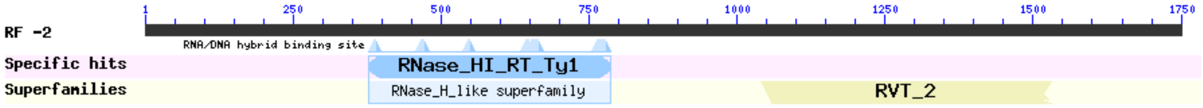


| chr0_nlr_300_Cc partial (pseudogene) | | | | |
|---|---|---|---|---|
| Name | Accession | Description | Interval | E-value |
| RNase_HI_RT_Ty1 | cd09272 | Ty1/Copia family of RNase HI in long-term repeat retroelements; Ribonuclease H (RNase H) | 378-785 | 3.94e-67 |
| RVT_2 super family | cl06662 | Reverse transcriptase (RNA-dependent DNA polymerase) | 1041-1529 | 4.39e-47 |

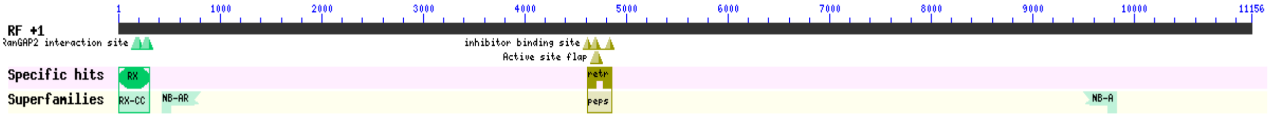


| chr8_nlr_26_Cc complete (pseudogene) | | | | |
|---|---|---|---|---|
| Name | Accession | Description | Interval | E-value |
| NB-ARC super family | cl26397 | NB-ARC domain | 427-801 | 1.88e-22 |
| NB-ARC super family | cl26397 | NB-ARC domain | 9502-9828 | 2.21e-15 |
| RX-CC_like | cd14798 | Coiled-coil domain of the potato virus X resistance protein and similar proteins | 4-300 | 2.90e-11 |
| retropepsin_like | cd00303 | Retropepsins; pepsin-like aspartate proteases; The family includes pepsin-like aspartate proteases from retroviruses, retrotransposons and retroelements | 4612-4851 | 3.61e-06 |

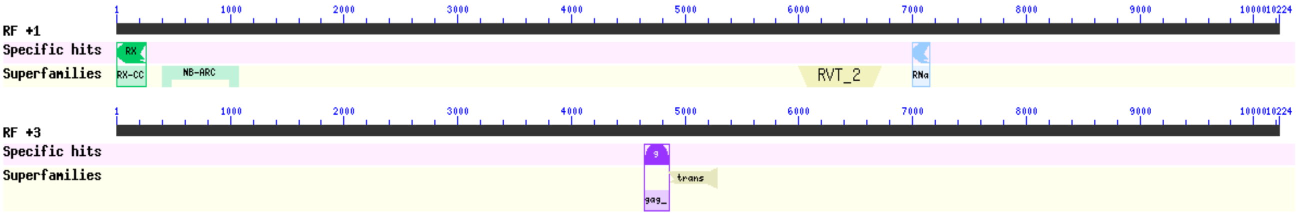


| chr7_nlr_33_Cc complete (pseudogene) | | | | |
|---|---|---|---|---|
| Name | Accession | Description | Interval | E-value |
| RVT_2 super family | cl06662 | Reverse transcriptase (RNA-dependent DNA polymerase) | 5992-6726 | 3.87e-67 |
| NB-ARC super family | cl26397 | NB-ARC domain | 403-1068 | 1.85e-50 |
| RNase_HI_RT_Ty1 | cd09272 | Ty1/Copia family of RNase HI in long-term repeat retroelements; Ribonuclease H (RNase H) | 7000-7149 | 1.94e-19 |
| RX-CC_like | cd14798 | Coiled-coil domain of the potato virus X resistance protein and similar proteins | 1-261 | 1.47e-11 |
| transpos_IS481 super family | cl41329 | IS481 family transposase | 4848-5276 | 1.29e-14 |
| gag_pre-integrs | pfam13976 | GAG-pre-integrase domain; This domain is found associated with retroviral insertion elements | 4638-4862 | 3.10e-12 |

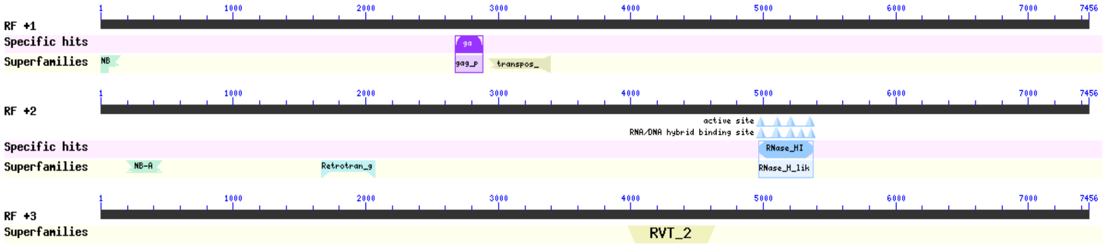

| chr4_nlr_22 Cc complete | | | | |
|---|---|---|---|---|
| Name | Accession | Description | Interval | E-value |
| transpos_IS481 super family | cl41329 | IS481 family transposase | 2935-3393 | 1.65e-13 |
| gag_pre-integrs | pfam13976 | GAG-pre-integrase domain; This domain is found associated with retroviral insertion elements | 2671-2886 | 4.35e-11 |
| NB-ARC super family | cl26397 | NB-ARC domain | 1-147 | 8.67e-07 |
| RNase_HI_RT_Ty1 | cd09272 | Ty1/Copia family of RNase HI in long-term repeat retroelements; Ribonuclease H | 4964-5377 | 1.25e-52 |
| Retrotran_gag_2 super family | cl26047 | gag-polypeptide of LTR copia-type; This family is found in Plants and fungi | 1664-2068 | 6.38e-22 |
| NB-ARC super family | cl26397 | NB-ARC domain | 194-457 | 4.89e-12 |
| RVT_2 super family | cl06662 | Reverse transcriptase (RNA-dependent DNA polymerase) | 3978-4631 | 7.19e-46 |

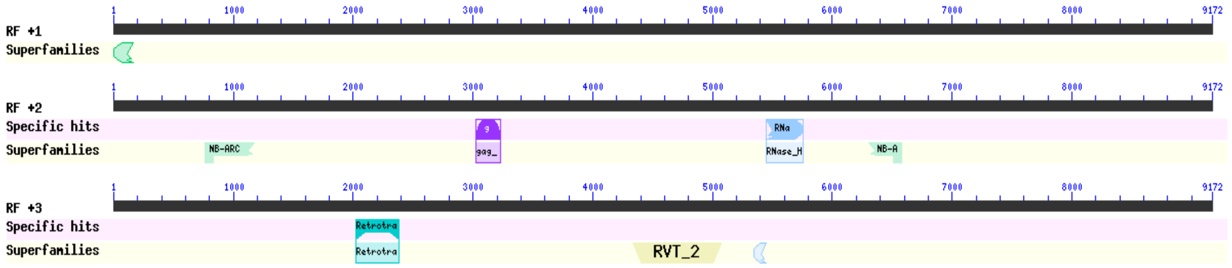


| **Chr_5_nlr_92_Ce complete** | | | | |
|---|---|---|---|---|
| **Name** | **Accession** | **Description** | **Interval** | **E-value** |
| RX-CC_like super family | cl36576 | Coiled-coil domain of the potato virus X resistance protein and similar proteins | 1-165 | 1.40e-07 |
| RNase_HI_RT_Ty1 | cd09272 | Ty1/Copia family of RNase HI in long-term repeat retroelements; Ribonuclease H | 5450-5752 | 1.79e-44 |
| NB-ARC super family | cl26397 | NB-ARC domain | 767-1171 | 1.40e-26 |
| gag_pre-integrs | pfam13976 | GAG-pre-integrase domain; This domain is found associated with retroviral insertion elements | 3026-3229 | 8.44e-13 |
| NB-ARC super family | cl26397 | NB-ARC domain | 6305-6580 | 5.76e-09 |
| RVT_2 super family | cl06662 | Reverse transcriptase (RNA-dependent DNA polymerase) | 4332-5072 | 3.19e-69 |
| Retrotran_gag_2 | pfam14223 | gag-polypeptide of LTR copia-type; This family is found in Plants and fungi | 2022-2381 | 3.58e-31 |
| RNase_H_like super family | cl14782 | Ribonuclease H-like superfamily, including RNase H, HI, HII, HIII, and RNase-like domain IV | 5343-5444 | 2.92e-08 |

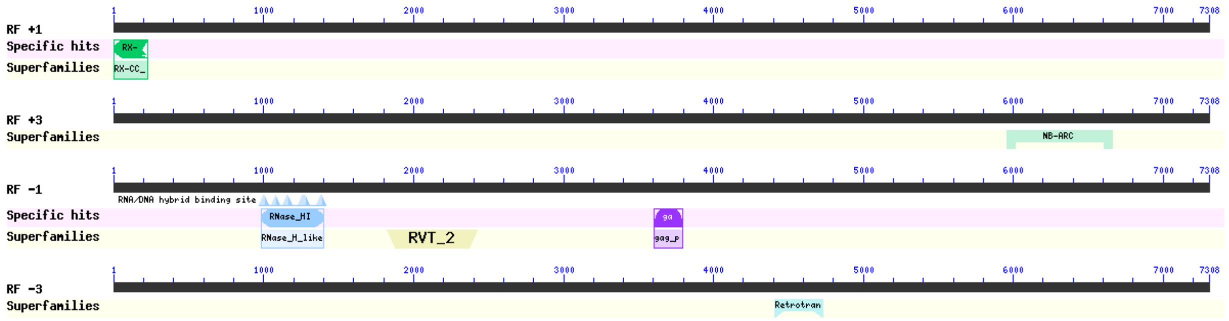


| **Chr_10_nlr_19_Ce complete (pseudogene)** | | | | |
|---|---|---|---|---|
| **Name** | **Accession** | **Description** | **Interval** | **E-value** |
| RX-CC_like | cd14798 | Coiled-coil domain of the potato virus X resistance protein and similar proteins; The potato | 1-225 | 8.90e-19 |
| NB-ARC super family | cl26397 | NB-ARC domain | 5955-6662 | 2.74e-43 |
| RNase_HI_RT_Ty1 | cd09272 | Ty1/Copia family of RNase HI in long-term repeat retroelements; Ribonuclease H | 986-1399 | 1.04e-62 |
| RVT_2 super family | cl06662 | Reverse transcriptase (RNA-dependent DNA polymerase) | 1823-2422 | 1.44e-40 |
| gag_pre-integrs | pfam13976 | GAG-pre-integrase domain; This domain is found associated with retroviral insertion elements | 3602-3796 | 2.47e-09 |
| Retrotran_gag_2 super family | cl26047 | gag-polypeptide of LTR copia-type; This family is found in Plants and fungi | 4407-4730 | 2.75e-10 |

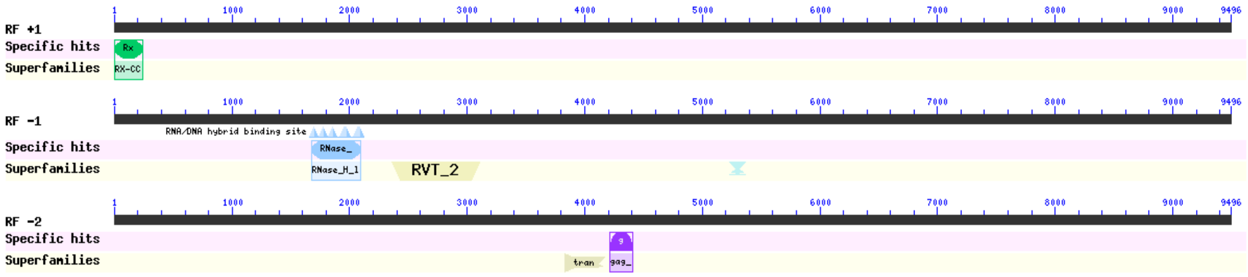


| **Chr_1_nlr_53_Ce complete (pseudogene)** | | | | |
|---|---|---|---|---|
| **Name** | **Accession** | **Description** | **Interval** | **E-value** |
| Rx_N | pfam18052 | This entry represents the N-terminal domain found in many plant resistance proteins | 1-243 | 8.01e-10 |
| RVT_2 super family | cl06662 | Reverse transcriptase (RNA-dependent DNA polymerase) | 2363-3100 | 2.71e-68 |
| RNase_HI_RT_Ty1 | cd09272 | Ty1/Copia family of RNase HI in long-term repeat retroelements; Ribonuclease H | 1676-2089 | 1.39e-64 |
| Retrotran_gag_2 super family | cl26047 | gag-polypeptide of LTR copia-type; This family is found in Plants and fungi | 5228-5365 | 4.61e-07 |
| transpos_IS481 super family | cl41329 | IS481 family transposase | 3826-4164 | 1.06e-15 |
| gag_pre-integrs | pfam13976 | GAG-pre-integrase domain; This domain is found associated with retroviral insertion elements | 4216-4407 | 1.18e-12 |

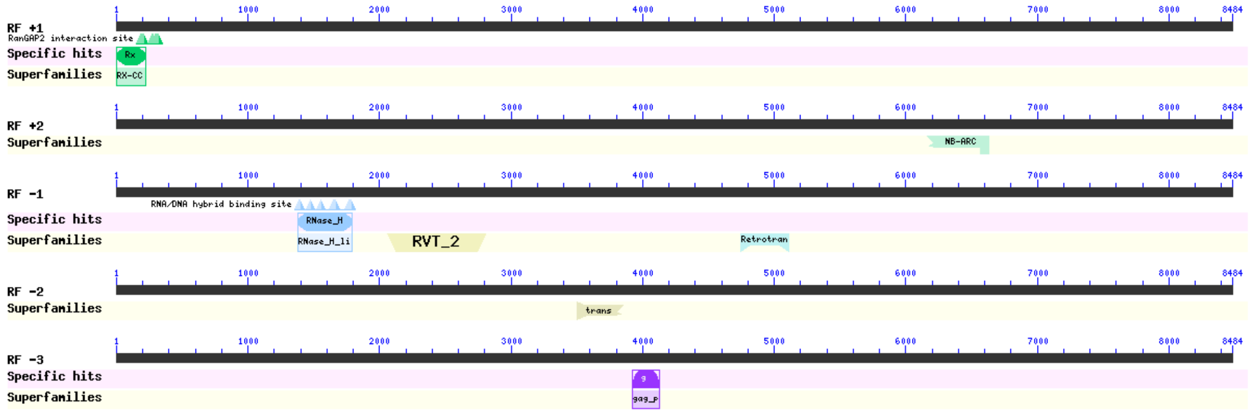


| Chr_8_nlr_53 complete (pseudogene) | | | | |
|---|---|---|---|---|
| Name | Accession | Description | Interval | E-value |
| Rx_N | pfam18052 | This entry represents the N-terminal domain found in many plant plant resistance proteins | 4-225 | 1.00e-16 |
| NB-ARC super family | cl26397 | NB-ARC domain | 6167-6631 | 6.51e-24 |
| RVT_2 super family | cl06662 | Reverse transcriptase (RNA-dependent DNA polymerase) | 2062-2802 | 1.81e-65 |
| RNase_HI_RT_Ty1 | cd09272 | Ty1/Copia family of RNase HI in long-term repeat retroelements; Ribonuclease H | 1381-1791 | 2.54e-57 |
| Retrotran_gag_2 super family | cl26047 | gag-polypeptide of LTR copia-type; This family is found in Plants and fungi | 4744-5106 | 2.87e-22 |
| transpos_IS481 super family | cl41329 | IS481 family transposase; | 3507-3848 | 3.91e-10 |
| gag_pre-integrs | pfam13976 | GAG-pre-integrase domain; This domain is found associated with retroviral insertion elements | 3923-4126 | 4.06e-13 |

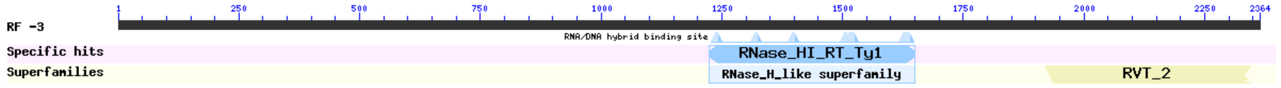


| Chr_8_nlr_67_Ce partial (pseudogene) | | | | |
|---|---|---|---|---|
| **Name** | **Accession** | **Description** | **Interval** | **E-value** |
| RNase_HI_RT_Ty1 | cd09272 | Ty1/Copia family of RNase HI in long-term repeat retroelements; Ribonuclease H (RNase H) | 1224-1649 | 7.59e-41 |
| RVT_2 super family | cl06662 | Reverse transcriptase (RNA-dependent DNA polymerase) | 1920-2345 | 8.89e-26 |

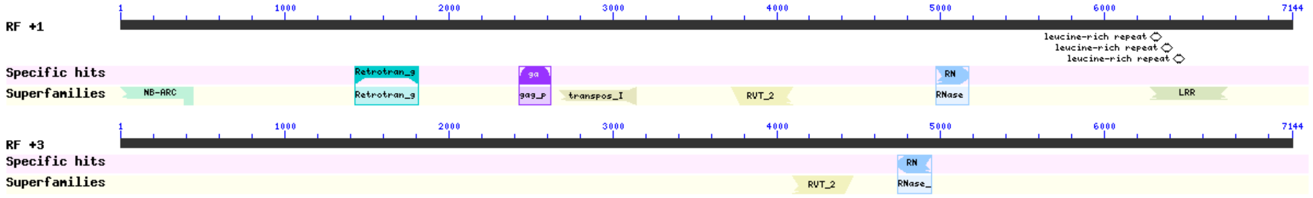


| Chr_11_nlr_129_Ce partial (pseudogene) | | | | |
|---|---|---|---|---|
| **Name** | **Accession** | **Description** | **Interval** | **E-value** |
| RVT_2 super family | cl06662 | Reverse transcriptase (RNA-dependent DNA polymerase) | 3721-4095 | 1.01e-43 |
| Retrotran_gag_2 | pfam14223 | gag-polypeptide of LTR copia-type; This family is found in Plants and fungi | 1429-1815 | 2.51e-29 |
| NB-ARC super family | cl26397 | NB-ARC domain | 1-438 | 5.97e-26 |
| RNase_HI_RT_Ty1 | cd09272 | Ty1/Copia family of RNase HI in long-term repeat retroelements; Ribonuclease H | 4972-5169 | 5.72e-23 |
| transpos_IS481 super family | cl41329 | IS481 family transposase | 2674-3144 | 4.64e-18 |
| gag_pre-integrs | pfam13976 | GAG-pre-integrase domain; This domain is found associated with retroviral insertion elements | 2431-2622 | 5.34e-10 |
| LRR super family | cl34836 | Leucine-rich repeat (LRR) protein [Transcription] | 6277-6747 | 2.28e-07 |
| RNase_HI_RT_Ty1 | cd09272 | Ty1/Copia family of RNase HI in long-term repeat retroelements; Ribonuclease H | 4737-4943 | 4.01e-30 |
| RVT_2 super family | cl06662 | Reverse transcriptase (RNA-dependent DNA polymerase) | 4095-4466 | 5.86e-23 |

**Supplementary Figure 2. Conserved domains analysis for loci from NLR-annotator that did not present homology to NLRs proteins by BLASTx analysis**. For the analysis, nucleotide sequences were used. The table shows the list of domain hits found for each of the sequences and the domains related to NLR proteins are highlighted in green. The conserved domains were detected using NCBI Conserved Domain Database and the graphical summary was set to concise view. If the alignment omitted more than 20% of the either the n- or c-terminus or both, the partial nature of the hit is indicated in the graphical display as domain with jagged edges. At the end of each ID, the source genome is identified, Ca: *C. arabica*, Cc: *C. eugenioides* and Ce: *C. eugenioides*. RF: reading frame.
