## Supplementary Text 1 for "Genome-wide identification, characterization, and comparative analysis of NLR resistance genes in *Coffea spp*"

**Supplementary Text 1. NLRs genes not found by the NLR-annotator in the coffee genomes**.

The overlap analysis made it possible to identify genes annotated in the reference genomes that did not overlap with any locus from NLR-annotator. To examine these genes, an NLR-parser analysis was performed on this set. After this analysis, we observed genes that did not obtain motifs detectable by NLR-parser and therefore did not present annotations in NLR-annotator. These loci were considered as "not found" by the tool. For *C. arabica*, two genes were not found, the LOC113737176 (XP_027120243.1 disease resistance protein RGA2-like) and the LOC113735982 (XP_027118739.1 probable disease resistance protein At4g19060) and for *C. canephora*, five genes were not found: Cc02_g12220 (Putative disease resistance protein At4g19050), Cc03_g10360 (Hypothetical protein with PFAM:PF00931), Cc07_g18800 (Hypothetical protein with PFAM:PF00931), Cc00_g21910 (Putative NBS-coding resistance gene protein -Fragment), and Cc00_g35420 (Putative Disease resistance protein RGA2).

For *C. eugenioides*, two situations occurred: the LOC113766771 (XP_027166726.1- probable disease resistance protein At5g43730) and the LOC113766774 (XP_027166730.1 - disease resistance protein At4g27190-like) did not present motifs detectable by NLR-Parser and consequently, were not annotated by NLR-annotator. As well, two genes (LOC113766615 - putative disease resistance protein RGA4 and LOC113777141 - putative late blight resistance protein homolog R1B-17) had at least three consecutive motifs belonging to the NB-ARC domain and detectable by NLR-Paser (standard threshold) but were not annotated. For the latter situation, a manual search of the positions in the txt file (Supplementary Table 3) was undertaken to make sure that there were no errors in the overlap analysis. The locus in this interval (Chr3 start at 40629207 and ends at 40630439, Ch7 start at 40630439 and ends at 22112852) was not detected by the NLR-annotator, and these genes were also classified as "not found" (Supplementary Table 6).

To ensure that the genes that NLR-annotator did not found encode proteins with NB-ARC domains, in addition to the PfamScan analysis, a conserved domains analysis using the nucleotide sequence was performed (Supplementary Figure 1). The goal here was to use a strategy similar to that used by the NLR-annotator. For *C. arabica*, we observed that the two genes belong in the NB-ARC superfamily, but with incomplete domain and variation in domain length. This also occurred in *C. canephora*, except for one gene (Cc07_g18800) that did not present detectable domains, despite showing a small fragment of the NB-ARC with PfamScan analysis (highlighted in orange in Supplementary Table 4). Additionally, in this genome, the Cc00_g35420 gene presented an NB-ARC incomplete domain and contained an N-terminal with Rx_N (pfam18052), a domain found in many plant resistance proteins. For *C. eugenioides*, two genes (LOC113766615 and LOC113766771) presented NB-ARC incomplete domains of different lengths, and the LOC113766774 gene, in addition to presenting NB-ARC complete domain, contains leucine-rich repeat. The gene LOC113777141 has two NB-ARC fragments separated by a domain belonging to the retrotran gag two superfamilies, containing a gag-polypeptide of LTR (long terminal repeat), which is a known type of retrotransposon in plants.
